# Tumor vascularization and the nonlinear dynamics of metastatic cancer treated by oncolytic viral therapy

**DOI:** 10.64898/2026.10.19.760241

**Authors:** Tyler P. Evans, Max Martin

**Affiliations:** Department of Mathematics, Northwestern University, 633 Clark St., Evanston, 60208, IL, United States of America; Department of Mathematics, University of Utah, 201 Presidents’ Circle, Salt Lake City, 84112, UT, United States of America

**Keywords:** Oncolytic viral therapy, mathematical oncology, nonlinear dynamics, bifurcation analysis

## Abstract

Oncolytic viral therapy (OVT) is an emerging cancer treatment where therapeutic viruses are administered to a patient with the goal of selectively targeting tumor cells while leaving healthy cells unaffected. Models have been developed which describe the interactions of the virus, tumor and immune system with systems of nonlinear ordinary differential equations (ODEs). Some models suggest that an initial phase of treatment with immunosuppressants could improve patient outcomes by preventing the immune system from attacking the oncolytic viruses before they can infect many tumor cells. However, all such models have focused on a single, localized tumor. Here, we model a primary tumor and its metastatic seeding of a secondary organ under simplifying assumptions about tumor vascularization. Bifurcation analysis and simulations reveal regions in parameter space for which immune modulation may encourage distant metastasis, leading to the dangerous scenario where the elimination of a primary tumor creates the false impression of a good prognosis. We also suggest that vascular disrupting agents may be useful for treating small tumors while they may promote OVT resistance in larger tumors.

## 1 Introduction

In recent years, oncolytic viral therapy (OVT) has emerged as a promising new treatment for cancer [1–4]. Viruses that selectively infect tumor cells, whether naturally occurring [5–7] or genetically engineered [8–10], are administered to patients. This promotes tumor destruction by two means: first, by directly lysing infected tumor cells and, second, by increasing the visibility of tumor cells to the immune system. When the immune system detects tumor cells, it readily attacks and destroys them [11, 12]; natural killer (NK) cells in particular exhibit strong anti-tumor [11, 12] activity. However, many clinically significant tumors contain adaptations that make them immunologically “cold” — that is, less likely to elicit a response from the immune system [13–15]. Once a tumor cell is infected, the patient’s immune system is more easily able to identify and attack it [16]. This approach has seen clinical successes [17–19]. However, the immune system also acts against the therapeutic virus [11, 12, 20], undermining the success of OVT. This leads to a complex, three-way interaction between the immune system, the tumor, and the virus [2, 21].

These interactions have been modeled and analyzed mathematically [22, 23], often using systems of nonlinear ordinary differential equations (ODEs) [21, 24–32]. Although simplified idealizations, these mathematical models have produced qualitative and even semi-quantitative insights of clinical importance. Of particular interest to the present work, [31, 33] have studied a model of immune-viral-tumor interactions for a single tumor, revealing conditions for, e.g., treatment failure. A major implication of this work, and related work [20, 32, 34], is the suggestion that an initial phase of treatment by immunosuppressive drugs may act to improve the therapeutic efficacy of OVT by preventing the immune system from destroying the therapeutic virus, ultimately promoting a greater level of viral buildup and tumor clearance.

However, the role of the immune system in the dynamics of metastasis is extremely complex and, in some cases, poorly understood [35, 36]. In recent mathematical models of OVT, metastatic dynamics also appear to be entirely absent; we imagine that this is due to the great deal of mathematical difficulty brought on by including them. At the same time, metastatic cancer is responsible for the majority of cancer deaths [37, 38], and the exact mechanisms of metastasis remain difficult to study. Of particular importance is explaining the apparent affinity of tumors originating in one organ for other specific organs. In view of the role that the immune system plays in keeping micrometastases dormant [16, 27, 39, 40], we ask whether adding immune suppressants to OVT might successfully destroy or suppress a primary tumor — or localized collection of oligometastases — only to inadvertently contribute to eventual, distant metastasis elsewhere in the body, or the “awakening” of dormant micrometastases. Indeed, the exact conditions under which micrometastases remain dormant are also imperfectly understood [16, 39, 40]. Similar concerns about the use of immune suppressants alongside OVT have been acknowledged elsewhere [41].

To address these questions from a mathematical perspective, we generalize a model due to [31] to the case of a primary tumor that may exhibit a variety of immunetumor and viral-tumor functional responses. These functional responses represent the differences between well-vascularized tumors, where the therapeutic virus and immune system have comprehensive access to the tumor, or avascular tumors, where the virus and immune system have access only to the tumor cells near the surface. Indeed, it is well-known that tumor geometry and vascularization themselves can act either to facilitate or to impede viral and other therapies [42, 43]. We newly include in this model a secondary site that receives an ongoing supply of micrometastases, developing a 6-component dynamical system. We apply a semi-analytical technique to obtain all equilibria and characterize their local stability. This enables two complementary lines of investigation. First, we conduct a bifurcation analysis, studying the stability of steady states corresponding to desirable clinical outcomes (e.g., the tumor-free equilibrium). Second, we perform simulations of the full dynamical system under several idealized treatment protocols, including combinations of OVT, immune modulation, and vascular disrupting agents (VDAs).

Our most significant findings are as follows. First, we find that tumor vascularization is an important determining factor in the success or failure of OVT: well-vascularized tumors appear to be more amenable to treatment by OVT, while avascular or hypovascular tumors may require supraphysiological parameter values to achieve complete tumor clearance with OVT alone. Second, when the metastasis to a secondary organ is modeled, we find scenarios where a primary tumor is successfully cleared by OVT but the development of a secondary tumor is encouraged. This can occur because the primary tumor is able to successfully seed a secondary tumor before it is eliminated; because the secondary tumor is simply less vulnerable to OVT than the primary tumor; or because the initial immune suppression phase prevents the immune response that would normally prevent growth. Notably, all of these effects are missed when considering a model that considers only one kind of tumor vascularization and does not handle metastasis. Finally, we find that the combination of OVT and VDAs may be broadly contraindicated; it appears that devascularizing a tumor can unintentionally shelter it against the anti-tumor effects of OVT.

## 2 Model

The proposed benefits of treatment by an oncolytic virus are twofold. First, the virus directly lyses tumor cells. Second, infected tumor cells are more easily detected by the patient’s immune system: that is, tumor cells which were previously immunologically undetectable may become detectable. However, the body’s own immune system recognizes the oncolytic virus and mounts an immune response to it. A robust immune system may thereby reduce the efficacy of OVT by destroying the virus before it has time to infect enough tumor cells to have a meaningful therapeutic effect. At the same time, tumor cells themselves can induce immune modulators, either promoting or suppressing local immunological activity. The need to understand these complex, multilateral interactions has led to interest in developing theoretical models that incorporate the main effects.

Here, we follow the approach of [27, 28, 33], and particularly [31], who proposed a system of nonlinear ordinary differential equations (ODEs),

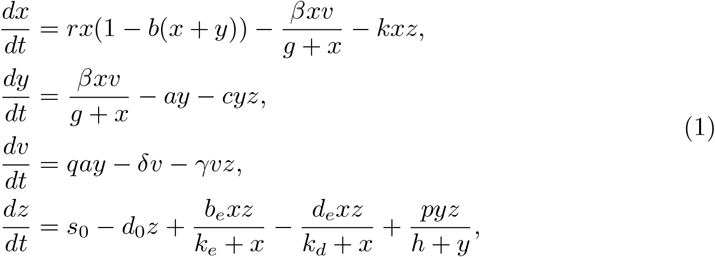

representing the dynamical interactions between immune cells (*z*), virions (*v*), tumor cells (*x*), and tumor cells that have been infected by the virus (*y*). Previous work has generally focused on the bifurcation structure and clinical implications of models of a single tumor and its multilateral interactions alongside the immune system and OVT [2, 27, 28, 31]. There has apparently been no work in this style which studies the possible interactions of tumors having multiple “phenotypes” — that is, where the immune-tumor or viral-tumor interactions are different across sites, such as a primary tumor and a nearby organ that might be the site of metastatic growth. This might occur, clinically, due to a solid, well-vascularized tumor shedding cells as avascular micrometastases [38, 44]. Although many tumors begin as avascular micrometastases before vascularizing, some, such as certain kinds of liver tumors [45], remain poorly vascularized even when large.

In view of the above discussion, we generalize Equations (1) in two ways: (i) the addition of a secondary organ that could form a new tumor, *x*_2_, which receives the micrometastases shed by the primary tumor *x*_1_; and (ii) the generalization of functional responses for tumor-virus and tumor-immune interactions. These considerations lead to the system of six ODEs,

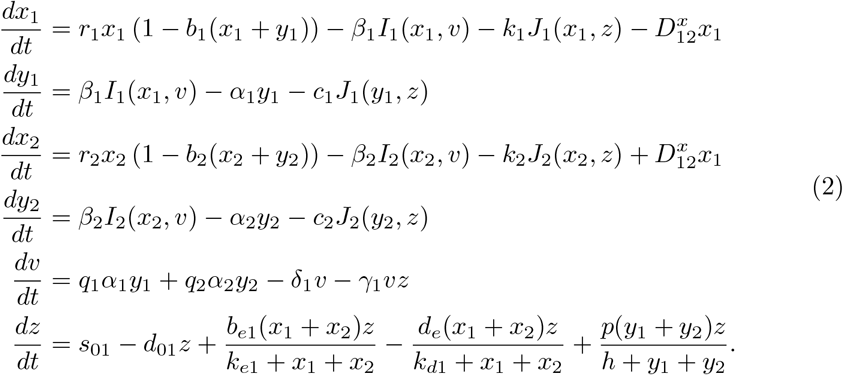

Above, logistic growth for tumor cells *x* with basic reproductive rate *r* and a carrying capacity *b*^*™*1^ is assumed. Both uninfected and infected tumor cells contribute to occupancy. Notably, we assume, as did [31], that *b*_1_ and *b*_2_ are fixed, static quantities. However, it is unclear whether the size of real tumors should be expected to saturate as a logistic growth law implies. An important implication of this is that, for realistic comparison between theory and clinical observations, parameter *b* may be unknowable *a priori*. Hence, in the present work, we will focus mostly on the qualitative dynamics of tumor growth as they depend on parameters that are more easily measured.

Tumor cells are infected at rate *βI*(*x, v*) by viral-tumor interactions, and eliminated entirely by immune-tumor interactions at rate *kJ*(*x, z*). Infected tumor cells *y* are created at rate *βI*(*x, v*), lysed at rate *αy*, and eliminated by immune interactions at rate *cJ* (*y, z*). Virions are contributed by the destruction of infected tumor cells *y* at rate *qαy*, are degraded naturally at rate *δv*, and destroyed by the immune system at rate *γvz*. The population of immune cells *z* recruits from the circulatory system at rate *s*_0_ and dies naturally at rate *d*_0_*z*. Additional immune cells are recruited at rate 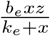due to interactions with uninfected tumor cells, while recruitment is also suppressed at rate 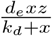 at rate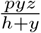. [13–15]. Recruitment due to interactions with infected tumor cells occurs

To consider differences in tumor vascularization, and as an important departure from previous work [31], we allow for general viral-tumor functional response, *I*(*x, v*), and immune-tumor functional response, *J*(*x, z*) or *J*(*y, z*). These interactions of tumor cells with virions and immune cells have been modeled variously throughout the literature. In particular, when the equations for *x*_2_ and *y*_2_ are ignored and one assigns the Michaelis-Menten functional response 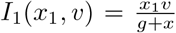, *J*_1_(*x*_1_, *z*) = *x*_1_*z*, and the Law of Mass Action functional response *J*_1_(*y*_1_, *z*) = *y*_1_*z*, Equations (2) specialize to Equations (1), i.e., those of Vithanage et al. [31]. In the present work, we will use a Michaelis-Menten response to model crowding effects due to low vascularization of the tumor mass, leading to poor mixing of immune cells and virions with tumor cells. A LOMA-like response is called for when immune cells and virions have unimpeded access to tumor cells due to a well-vascularized tumor with no rate limits. Other functional response models have been considered elsewhere; for example, Beddington-deAngelis functional responses have been used [27], as have ratio-dependent responses informed by cellular automata models [46].

To summarize, functional responses *I* and *J* can each take on two different forms:

- Law of Mass Action forms, *I*(*x, v*) = *bxv, J*(*x, z*) = *bxz* and *J*(*y, z*) = *byz*; and
- Michaelis-Menten forms, 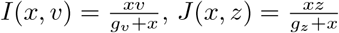, and 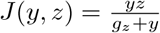, where the extra factor of *b* is used to ensure that the Law of Mass Action and Michaelis-Menten functional responses are dimensionally consistent. This ultimately simplifies the model nondimensionalization of Section 3.

Next, towards having a tractable — even if idealized — mathematical model, we further assume that tumor cells are shed from the primary tumor (site 1) to secondary organs (site 2) linearly at a rate of roughly one percent of tumor mass per day [47, 48], thereby introducing a source term in the equation for *x*_2_ of the form 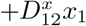. The assumption of linear shedding is a rather strong one. However, while an imperfect proxy for circulating tumor cells (CTC), which play an important role in distant metastasis [49, 50], shedding of ctDNA at early stages is, indeed, approximately linear with tumor size [48]. For larger, more developed tumors, physical limitations lead to sublinear scaling in the rate at which CTC arrive at a potential colonization site [51–53]. This is consistent with observations of T241 fibrosarcoma development in mice [47], where early exponential growth eventually saturates. Among the physical limitations contributing to growth rate saturation are intravasation difficulties and high shear stresses experienced by tumor cells in the bloodstream [52–55]. Moreover, internal stresses generated by tumor growth can destroy the tumor’s own vasculature [56–59]. Since CTC shedding is a perfusion-dependent process [54], this implies that the linear relationship cannot persist. Indeed, tumor vascular disrupting agents (VDAs) have gained attention recently for their potential therapeutic use in stifling the metastatic cascade [60]. Taken together, this broadly suggests that a Michaelis-Menten-like law should also apply to the shedding rate — that is, approximately linear shedding for *x*_*i*_ ≪ *m*_*i*_, with saturation for *x*_*i*_ *≫m*_*i*_, where *m*_*i*_ is a critical tumor mass threshold. The use of Michaelis-Menten-like dynamics for tumor cell proliferation is also suggested in [61]. However, the need to approximate reasonable values of *m*_*i*_ introduces into the present work additional, difficult-to-estimate parameters and increases computational complexity. We instead note that incorporation of these physical limitations is a worthwhile consideration for future work, while also unlikely to reverse our main findings.

As a further modeling simplification, we do not explicitly model the effects of cross-seeding of the primary tumor and second tumor; our assumption is equivalent to supposing that the primary tumor seeds to the secondary tumor, but not vice-versa. Although successful reverse-colonization of a primary tumor by its metastases is rare, tumor self-seeding and cross-seeding are increasingly recognized as important effects in metastasis [62–66]. We have also phenomenologically modeled all immune cells as the single quantity *z* without making any distinction between, e.g., lymphocytes, NK cells, etc., as has been done elsewhere [27]. Moreover, we make no distinction between immune cells near the primary tumor and those that are near the secondary site. This serves to reduce the algebraic difficulty of finding steady states (see Section 3), while also adding to the model the physical assumption that either the primary and secondary sites are not far away, or that the immune system is well-mixed at all times. A similar well-mixedness assumption is made for *v*. In a more detailed model, immune and virion transport might be better modeled as their own Michaelis-Menten-like terms, with saturation parameters tuned to distinguish between tissue types. Indeed, virions, tumor cells and infected tumor cells all appear to contribute to immune cell chemotaxis [67–70].

Hence Equations (2) are intended as an idealized, minimal model of a primary tumor and its metastasis that also respects recent advances in understanding the non-linear dynamics of OVT. It is sufficient for our present purposes, but should ultimately be supplanted with a more detailed spatiotemporal model. Finally, we comment that [31] proved the well-posedness of Equations (1). It is natural to suppose, then, that Equations (2) are also well-posed, being a generalization of [31]; this turns out to be the case, and the proof follows the same style as that of [31]. In Table 1, we record the baseline parameters used in [31], where they were broadly validated against a model of human pancreatic cancer in mice [71]. These same parameters will be used in the present work for comparison, along with an estimate for the shedding rate of the pri-mary tumor 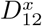 of around 0.1% to 1% of tumor cells per day [47, 72]; however, we note that this parameter assignment is complicated by a lack of first-principles modeling of *in vivo* tumorigenicity [73]. We also either record a plausible range or state that we have fixed the parameter as an assumption.

**Table 1.**
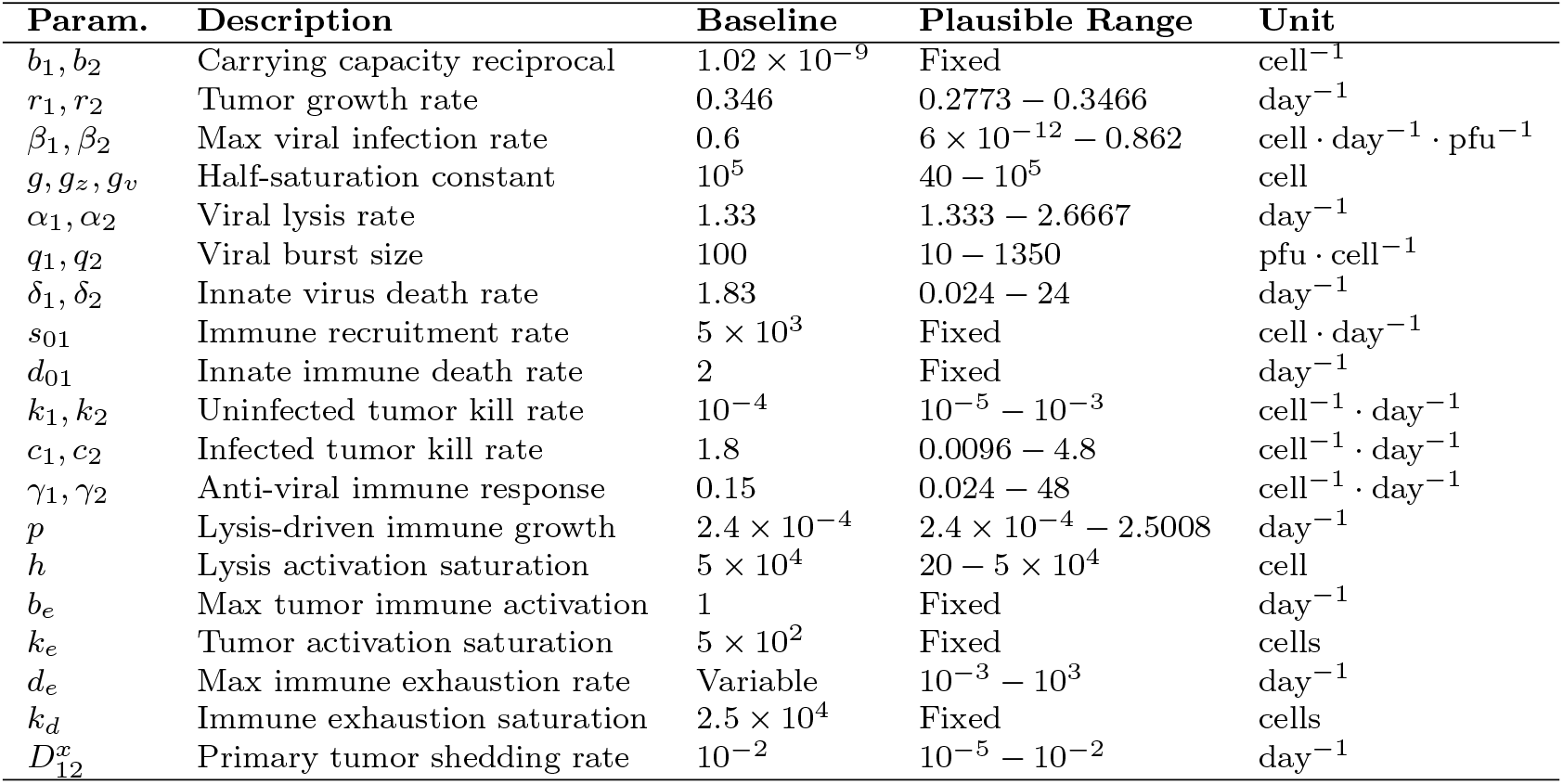
Model Parameters and Plausible Ranges.

| Param. | Description | Baseline | Plausible Range | Unit |
|---|---|---|---|---|
| b<sub>1</sub>, b<sub>2</sub> | Carrying capacity reciprocal | 1.02 × 10<sup>−9</sup> | Fixed | cell<sup>−1</sup> |
| r<sub>1</sub>, r<sub>2</sub> | Tumor growth rate | 0.346 | 0.2773 − 0.3466 | day<sup>−1</sup> |
| β<sub>1</sub>, β<sub>2</sub> | Max viral infection rate | 0.6 | 6 × 10<sup>−12</sup> − 0.862 | cell · day<sup>−1</sup> · pfu<sup>−1</sup> |
| g, g<sub>z</sub>, g<sub>v</sub> | Half-saturation constant | 10<sup>5</sup> | 40 − 10<sup>5</sup> | cell |
| α<sub>1</sub>, α<sub>2</sub> | Viral lysis rate | 1.33 | 1.333 − 2.6667 | day<sup>−1</sup> |
| q<sub>1</sub>, q<sub>2</sub> | Viral burst size | 100 | 10 − 1350 | pfu · cell<sup>−1</sup> |
| δ<sub>1</sub>, δ<sub>2</sub> | Innate virus death rate | 1.83 | 0.024 − 24 | day<sup>−1</sup> |
| s<sub>01</sub> | Immune recruitment rate | 5 × 10<sup>3</sup> | Fixed | cell · day<sup>−1</sup> |
| d<sub>01</sub> | Innate immune death rate | 2 | Fixed | day<sup>−1</sup> |
| k<sub>1</sub>, k<sub>2</sub> | Uninfected tumor kill rate | 10<sup>−4</sup> | 10<sup>−5</sup> − 10<sup>−3</sup> | cell<sup>−1</sup> · day<sup>−1</sup> |
| c<sub>1</sub>, c<sub>2</sub> | Infected tumor kill rate | 1.8 | 0.0096 − 4.8 | cell<sup>−1</sup> · day<sup>−1</sup> |
| γ<sub>1</sub>, γ<sub>2</sub> | Anti-viral immune response | 0.15 | 0.024 − 48 | cell<sup>−1</sup> · day<sup>−1</sup> |
| p | Lysis-driven immune growth | 2.4 × 10<sup>−4</sup> | 2.4 × 10<sup>−4</sup> − 2.5008 | day<sup>−1</sup> |
| h | Lysis activation saturation | 5 × 10<sup>4</sup> | 20 − 5 × 10<sup>4</sup> | cell |
| b<sub>e</sub> | Max tumor immune activation | 1 | Fixed | day<sup>−1</sup> |
| k<sub>e</sub> | Tumor activation saturation | 5 × 10<sup>2</sup> | Fixed | cells |
| d<sub>e</sub> | Max immune exhaustion rate | Variable | 10<sup>−3</sup> − 10<sup>3</sup> | day<sup>−1</sup> |
| k<sub>d</sub> | Immune exhaustion saturation | 2.5 × 10<sup>4</sup> | Fixed | cells |
| D<sup>x</sup><sub>12</sub> | Primary tumor shedding rate | 10<sup>−2</sup> | 10<sup>−5</sup> − 10<sup>−2</sup> | day<sup>−1</sup> |

## 3 Analysis

### 3.1 Model nondimensionalization

Models having as many parameters as Equations (2) commonly become difficult to analyze. In addition, parameters can be difficult to obtain experimentally, or to estimate. They might also be known only up to a range of uncertainty. To address these difficulties, we nondimensionalize Equations (2), introducing the dimensionless variables

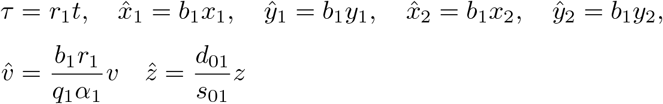

and assigning primacy to the main tumor’s parameters. This nondimensionalization of Equations (2) leads one to encounter the dimensionless parameter groups

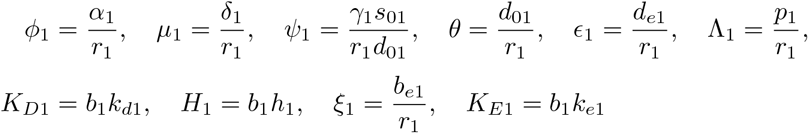

which occur regardless of functional responses *I* and *J*. We also encounter the dimensionless parameter groups

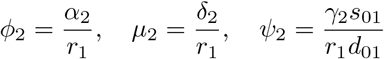

which are associated with the secondary site, and dimensionless parameter groups

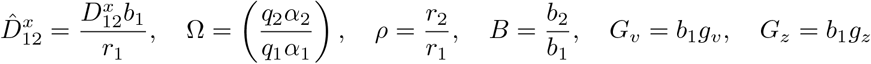

which relate the two sites. We also encounter the quantities

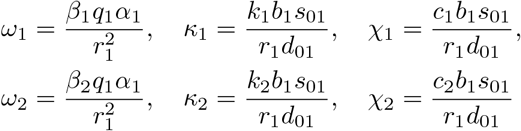

which, in contrast with the others, *are not yet dimensionless*; rather, they each have units cell^*™*1^. Here, one encounters two interconnected issues:

- Given that the Law of Mass Action and Michaels-Menten functional forms involve different parameters (*b* for Law of Mass Action and *g*_*z*_ or *g*_*v*_ for Michaelis-Menten), how should the nondimensionalization accommodate this?
- How can a fair, quantitative comparison be made between functional response types with a shared set of parameters (Table 1) that were validated for only one specific functional response combination, *I*_*MM*_, *J*_*LOMA*_?

To simultaneously address these two points, we observe that the parameters of Table 1 were validated for an *I*_*MM*_, *J*_*LOMA*_-type tumor [31]. In principle, the present model could be applied to the same experimental data as [31]; however, this would fundamentally alter the nature and intention of the present work. Instead, and with the intention of preserving the physical meaning of Table 1’s parameters while ensuring that they are fully dimensionless, we apply the following transformations. For a tumor having infectivity parameter *ω* and immune killing parameters *κ* and *χ*, we rescale as

- *κ → κ/b, χ → χ/b* when considering a tumor of phenotype *I*_*MM*_, *J*_*LOMA*_;
- *ω → ω/*(*gb*), *κ → κ/b, χ → χ/b* when considering a tumor of phenotype *I*_*LOMA*_, *J*_*LOMA*_; and
- *κ → κg*_*z*_, *χ → χg*_*z*_ when considering a tumor of phenotype *I*_*MM*_, *J*_*MM*_ .

We note that in all cases the division by *b* or multiplication by *g*_*z*_ complete the nondimensionalization. The above procedure also ensures that functional responses for low concentrations agree across all phenotypes. The difference between phenotypes only becomes apparent for higher concentrations. In the Appendix, we show algebraically that this nondimensionalization scheme properly recovers the intended dimensional quantities.

The parameter groups shown above have natural interpretations: 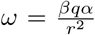 is the *effective viral infection rate*; 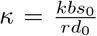 is the *immune killing rate of uninfected cells*;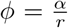is the *infected cell lysis rate relative to tumor growth*; 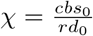is the *immunekilling rate of infected cells*; 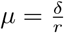 is the *viral clearance rate*; 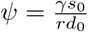 is the *immune-mediated viral clearance rate*; 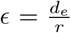 is the *baseline immune turnover rate*; 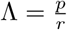 is the *tumor-induced immunosuppression rate*; 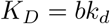 is the *infected-cell induced immunerecruitment rate*; *K*_*D*_ = *bk*_*d*_ and *H* = *bh* are *rescaled saturation constants*.

This nondimensionalization converts Equations (2) to

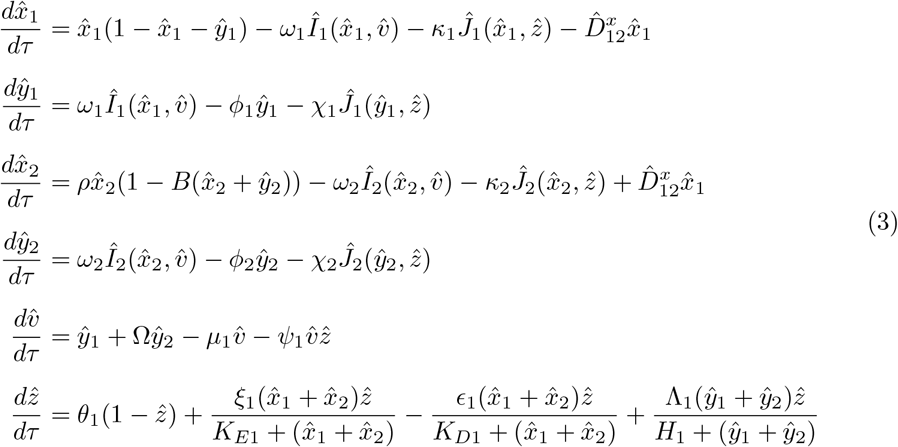

and reduces the number of free parameters for Equations (2), while also facilitating exploration of model predictions. In what follows, we will denote the functional forms

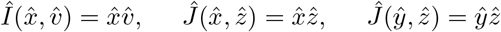

with a subscript “LOMA” for “Law of Mass Action” functional response (*I*_*LOMA*_ or *J*_*LOMA*_), and

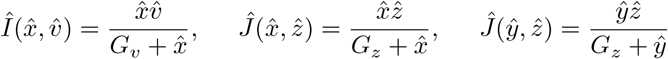

with a subscript “MM”, for “Michaelis-Menten” functional response (*I*_*MM*_ or *J*_*MM*_ ).

### 3.2 Equilibria and stability

Mathematically, a major goal of the present work is to develop a bifurcation analysis of Equations (2) with an eye towards understanding the dependence of the bifurcation structure on immune-tumor functional response and viral-tumor functional response. In general, a bifurcation analysis consists of three parts: (i) identifying the equilibria of a given dynamical system, (ii) determining the eigenvalues of the system’s Jacobian when evaluated at each of the equilibria, and (iii) sweeping across parameter space to identify changes in equilibria and stability.

Determining all equilibria of systems such as Equations (3) is exceedingly difficult. Here, we adopt a simple semi-analytical method. First, as is typical of a bifurcation analysis, we set all derivatives with respect to nondimensionalized time *τ* occurring in Equations (3) to zero. That is, we consider the nonlinear algebraic system

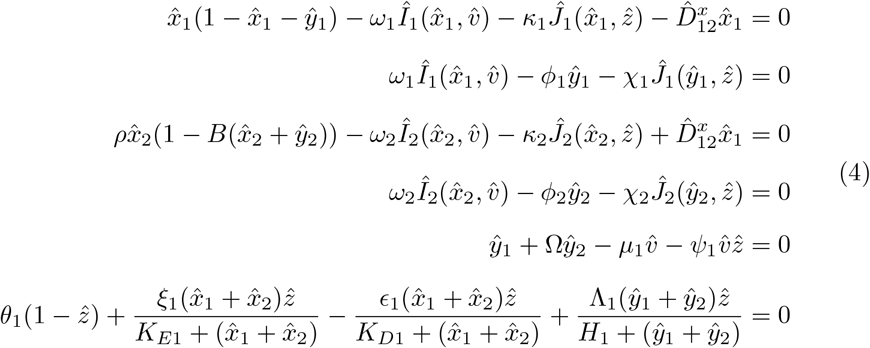

whose solutions 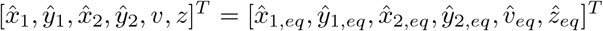 are said to be *equilibria* of Equations (3). In general, one cannot determine all of the solutions by hand, although some can be found. To simplify the problem of finding equilibria, we reduce Equations (4) to either a single polynomial in one variable, or a pair of mixed polynomials in two variables. This process is described as follows.

#### Reduction to polynomial equations

Our bifurcation analysis is concerned with two cases: (i) the scenario where we consider the primary tumor only and ignore the possible secondary tumor, setting 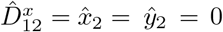, corresponding to Equations (9) of Section 4.1; and (ii) the scenario where the primary tumor has been eliminated 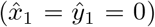 but the secondary tumor has not *necessarily* been eliminated, corresponding to Equations (10) of Section 4.2. In both cases, the strategy for reduction to polynomial equations is as follows. First, we observe that the equilibria are computed from a system of only four algebraic equations; dropping subscripts, these equations are of the form

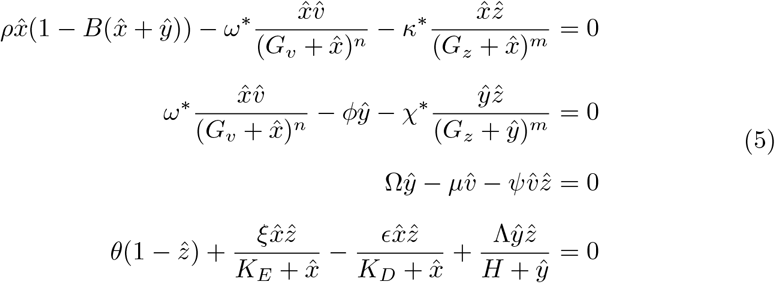

where *n, m* ∈ *{*0, 1} depend on which functional response is used, and up to the assignment of parameters *ρ, B*, and Ω. For Equations (5), there are three families of equilibria:

- 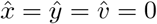, 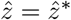, the equilibrium in which the tumor has been eliminated;
- 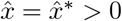, *ŷ*= 0, 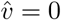, 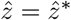, the equilibrium in which OVT has failed; and
- 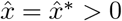, *ŷ*= *ŷ*^*\**^ > 0, 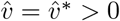, 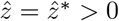, the equilibrium in which the tumor, immune system, and virion populations reach a “stalemate”.

In the first scenario, one immediately finds that Equations (5) reduce simply to

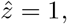

and the resulting equilibrium 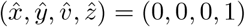 regardless of functional responses *I* and *J*. In the second, only Equations (5.1) and (5.4) remain. With the assumption that 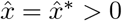, they have the form

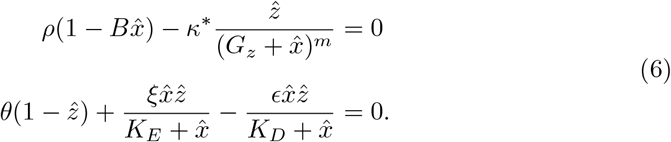

Equation (6.1) is solved for 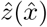. This is substituted into Equation (6.2) in order to obtain an algebraic equation completely in terms of 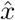. Multiplying through by 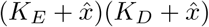, it is fully reduced to a single-variable polynomial equation for 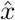.

In the third scenario, we again have 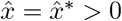, leading to

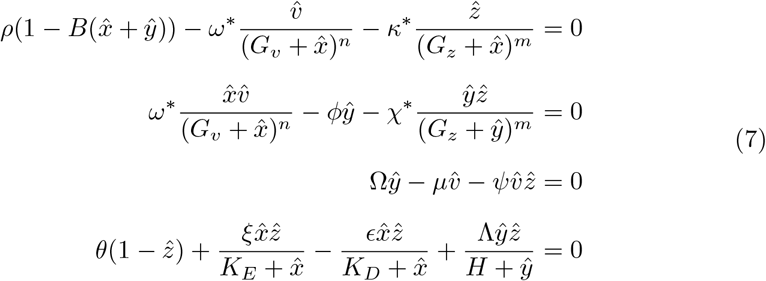

Here, we obtain a pair of polynomial equations in the variables 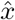 and 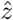 by solving for 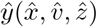 from Equation (7.1), substituting this into Equations (7.2)-(7.4). Then 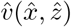 is isolated from Equation (7.2) and substituted into Equations (7.3)-(7.4). After clearing the denominators, this produces a pair of high-order polynomials in 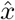 and 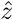.

#### Linearized dynamics

Once these polynomials are obtained, they are passed to Matlab’s *vpasolve*, which is efficient at solving small systems of polynomial equations. Next, we substitute the usual linear stability ansatz,

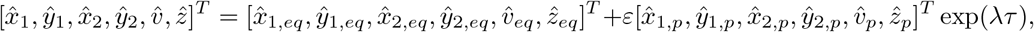

to Equations (3), where it is understood that *ε* ≪ 1. Taylor expansion up to *O* (*ε*) then leads to the eigenvalue problem

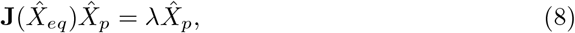

for eigenpair 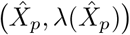, where 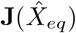 is the Jacobian of Equations (3) evaluated at equilibrium

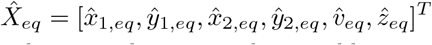

and we similarly denote the perturbation to this equilibrium as

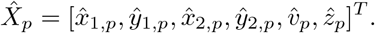

The Jacobian matrix **J** = (*J*_*i,j*_) ∈ ℝ^6×6^ is recorded in the Appendix.

For a fixed set of system parameters, we solve Equations (4) for 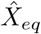 using *vpasolve* followed by computing the eigenvalues *λ* of 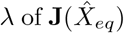, again in Matlab. We restrict atten-tion to physical solutions; that is, we consider only 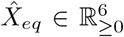 . Since 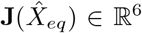,we obtain six eigenvalues *λ* ∈ C (counting multiplicity) for each equilibrium 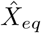. For a given equilibrium, if Re(*λ*) > 0 for any *λ*, the equilibrium is considered *unstable*. If, on the other hand, Re(*λ*) < 0 for all *λ*, the equilibrium is considered to be *stable*. For the present purposes, this simple classification is sufficient: we need not make distinc-tions between, e.g., spirals, saddles, etc., nor their higher-dimensional counterparts.

Moreover, the numerical simulations of Section 5.6 provide some visual representation of whether the long-term behavior suggested by the bifurcation structure is actually realized.

## 4 Results: bifurcation analysis

In this Section, we will summarize our main bifurcation results. First, we consider Equations (3) with the secondary site equations deleted, and the influence of functional responses *I* and *J* on the bifurcation structure of equilibrium values of 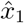as immune exhaustion parameter *d*_*e*_ and basic immune killing rate *k* are adjusted. As part of this, we reproduce some of the bifurcation diagrams of [31] as a special case.

Second, we introduce the two-site model, assume that the primary tumor has been eliminated 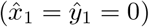, and examine the bifurcation structure of equilibrium values of 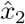 as *d*_*e*_ and *k* are adjusted. This serves to address one of our primary questions of interest — whether OVT can destroy a primary tumor while inadvertently promoting metastasis at a secondary site — while also making simple visualization possible. Moreover, we find that attempting to reduce the full six-component model to a small system of polynomials is prohibitively difficult, and numerical solutions are unwieldy.

### Note on parameter selection

In principle, Equations (3) allow for the use of different parameter values for the primary tumor and secondary organ. However, in order to maintain the relative simplicity of this work, we will assume that all parameters in Table 1 are shared between both tumors. These are the same parameter values used in [31] and validated there against a mouse model. This ensures that the results of the present work cannot be attributed to, e.g., having found an exceptional region in parameter space. It is also biologically reasonable: one often expects that the secondary tumor should be fairly similar to the primary tumor from which it originated, although there are notable exceptions [74–76].

### 4.1 Model with primary tumor only

We first focus on modeling only a primary tumor whose vascularization might be different from that considered in [31] and related work [28, 33]. In Figures 1-3, we present bifurcation diagrams for Equations (3) in the reduced case where the secondary site is ignored. That is, we set 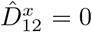, the equations for the evolution of *x*_2_ and *y*_2_ are deleted, and we set *x*_2_ = *y*_2_ = 0 elsewhere in the remaining equations. Specifically, we have the equations

**Fig. 1.**
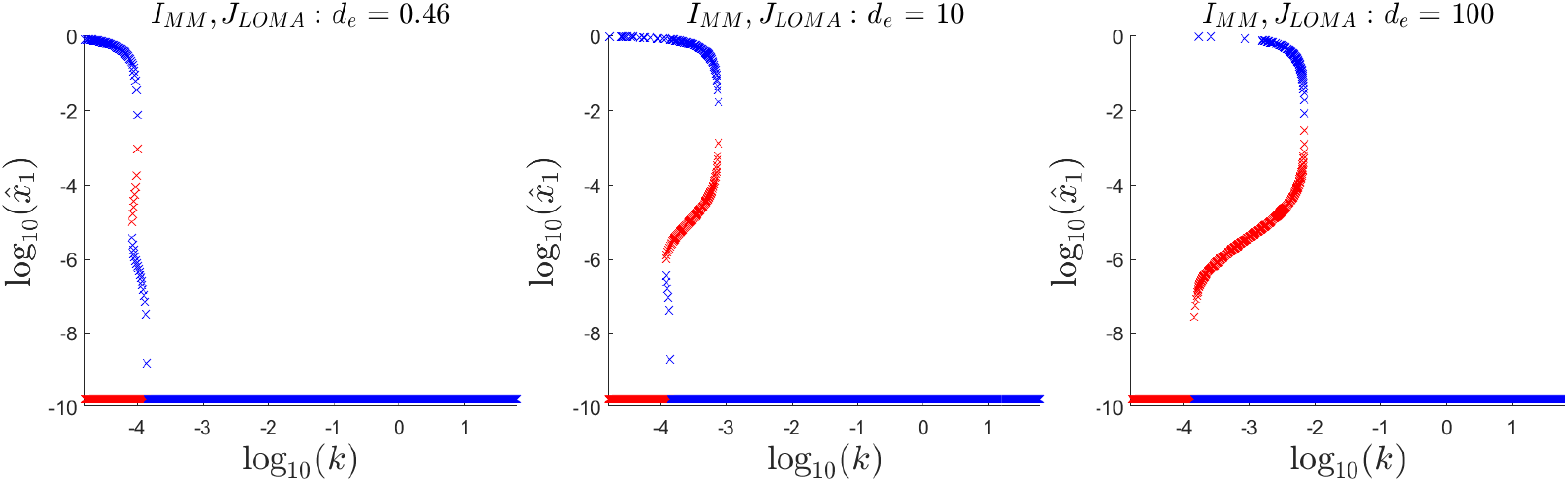
Bifurcation diagram for Equations (3) in the case of a primary tumor only where stable equilibrium values of 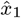 are colored blue and unstable values are colored red. Setting the infection rates *I* to Michaelis-Menten form and killing rates *J* to Law of Mass Action form specializes the present model to that of [31]. All parameters are as shown in Table 1. We also follow [31, 33] in superimposing the tumor-free state on the logarithmic scaling by assigning it to a value of log_10_ 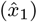 less than all others. This branch, nonetheless, corresponds to 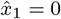.

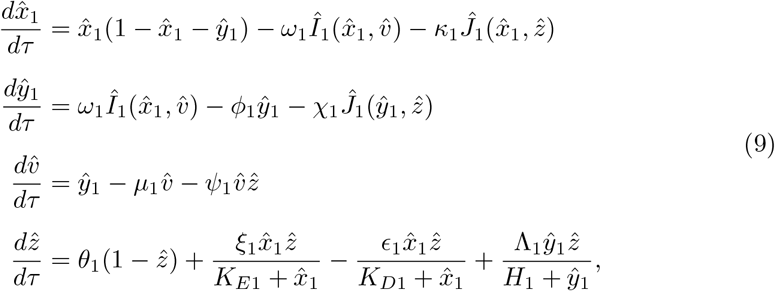

which constitute a single-site generalization of the model due to [31]. For several values of immune exhaustion parameter *d*_*e*_ and for each of three combinations of functional responses, equilibrium values of log_10_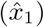 are plotted against the basic immune killing rate log_10_(*k*). When the equilibrium value is stable, the point is colored blue; when it is unstable, the point is colored red. The logarithmic scaling usefully emphasizes the shape of the bifurcation structure.

In particular, Figure 1 is essentially a reproduction of Figures 7-9 of [31] and serves partially as validation of the present work against previous work. As in [31], we find that increasing immune exhaustion *d*_*e*_ from 0.46 to 100 produces a transcritical bifurcation with a stable, small-tumor branch at low values of *d*_*e*_ becoming an unstable branch. The observation of [31] is that this unstable branch can be used as an approximation for the separatrix in parameter space — that is, for fixed values of *k* and *d*_*e*_, initial tumor sizes below the unstable red branch are drawn into the stable, tumor-free equilibrium.

Figure 2 shows how the bifurcation structure changes when the infection rate of tumor cells by virions proceeds according to the Law of Mass Action instead of a Michaelis-Menten response. As in Figure 1, a lower, stable small-tumor branch undergoes a transcritical bifurcation as *d*_*e*_ is increased, becoming unstable. Notably, however,the upper, stable large-tumor branch is *also* destabilized, in contrast with Figure 1 where it remains stable for all *k* and *d*_*e*_. While one anticipates that the lower, desta-bilized branch still represents an approximation for the separatrix, the newly unstable large-tumor branch suggests that beyond some critical *d*_*e*_, a well-vascularized tumor whose size lies in between the two unstable branches undergoes oscillations associated with the formation of a limit cycle. The tumor size might not stabilize to a single value.

**Fig. 2.**
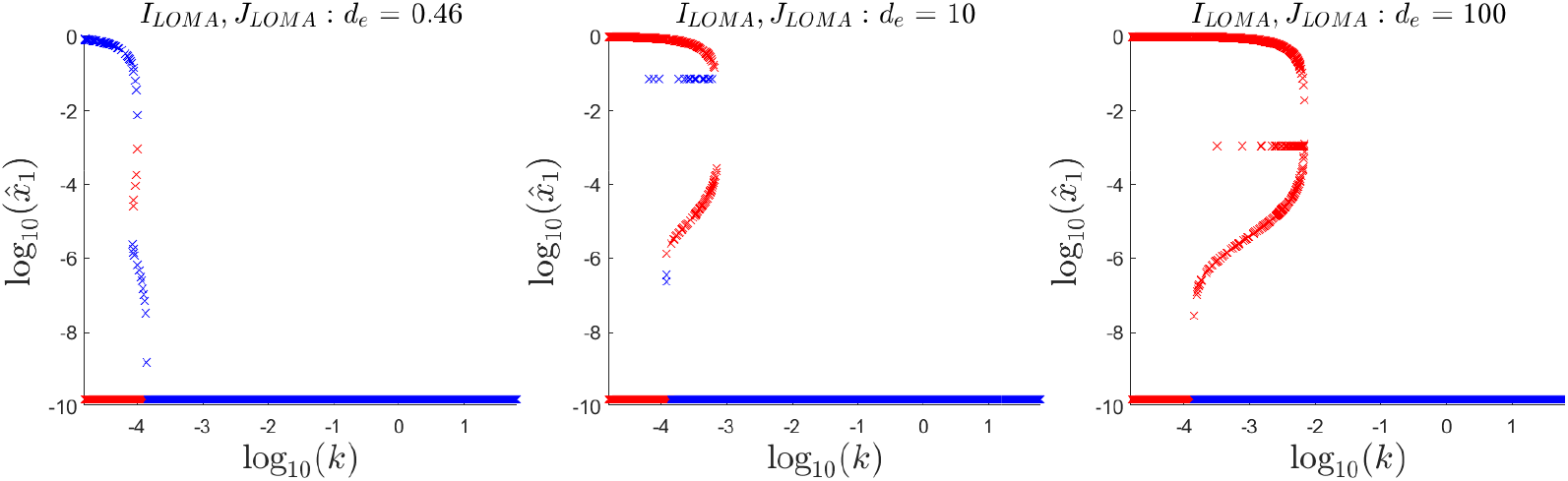
Bifurcation diagram for Equations (3) in the case of a primary tumor only where stable equilibrium values of 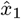 are colored blue and unstable values are colored red. Setting both the infection rates *I* and killing rates *J* to Law of Mass Action forms specializes the present model to the scenario of a well-vascularized tumor where the oncolytic virus and immune cells have unimpeded access to tumor cells. We also follow [31, 33] in superimposing the tumor-free state on the logarithmic scaling by assigning it to a value of log_10_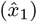less than all others. This branch, nonetheless, corresponds to 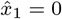.

In Figure 3, we consider the bifurcation diagrams for a primary tumor that is poorly vascularized. The infection rates *I* and immune killing rates *J* are both assumed to follow a Michaelis-Menten functional response. Here, the bifurcation structure is quite different to those shown in Figures 1 and 2. The transcritical bifurcation that destabilizes the stable small-tumor branch as *d*_*e*_ is increased is preserved. This unstable small-tumor branch is exceptionally long with respect to the *k* axis in comparison with those from *I*_*LOMA*_, *J*_*LOMA*_ and *I*_*MM*_, *J*_*LOMA*_ functional responses, suggesting that poorly vascularized small tumors are exceptionally treatable with OVT compared to those that are better vascularized. However, unlike Figures 1 and 2, the basic immune killing rate *k* needed so that only the stable, tumor-free equilibrium remains is at a supraphysiological level: there is no reasonable level of immune boosters that can be administered so that a tumor originating between the upper, large-tumor stable branch and the unstable small-tumor branch is destroyed. Hence Figure 3 represents a class of tumor that, when small, is easily suppressed; however, when it is only noticed at a late stage of development, it is especially difficult to treat using any combination of OVT and immune modulation.

**Fig. 3.**
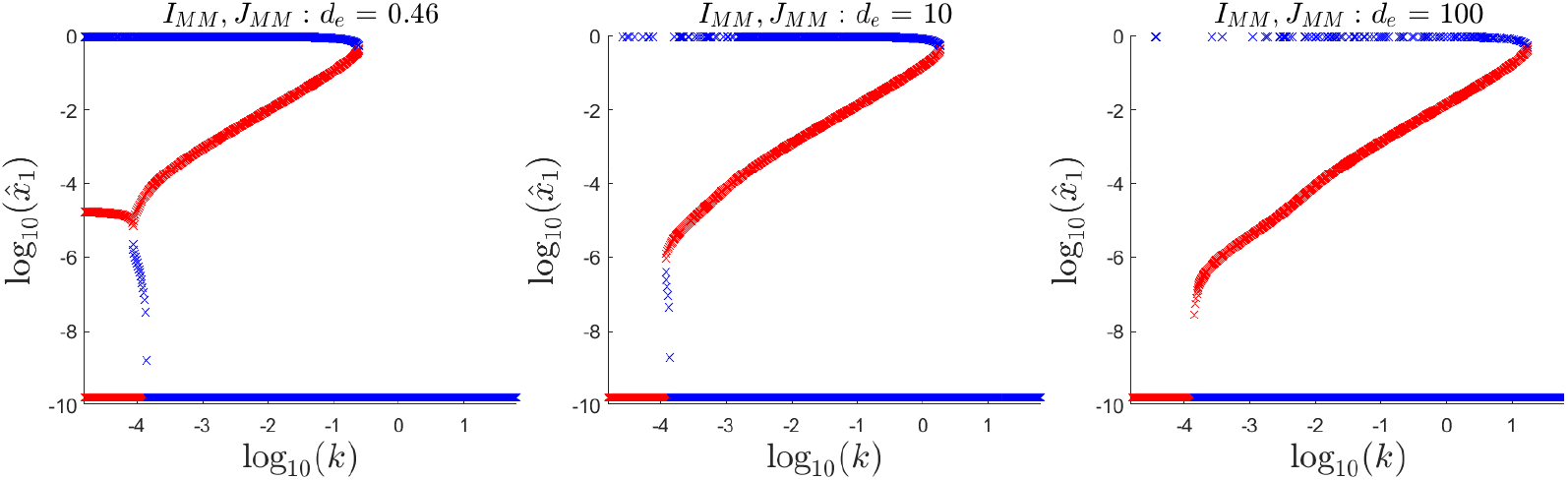
Bifurcation diagram for Equations (3) in the case of a primary tumor only where stable equilibrium values of 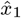are colored blue and unstable values are colored red. Setting both the infection rates *I* and killing rates *J* to Michaelis-Menten forms specializes the present model to the scenario of a poorly-vascularized tumor where the oncolytic virus and immune cells have only rate-limited access to tumor cells. We also follow [31, 33] in superimposing the tumor-free state on the logarithmic scaling by assigning it to a value of log_10_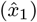 less than all others. This branch, nonetheless, corresponds to 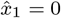.

### 4.2 Model with primary tumor and secondary site

Having studied models of the primary tumor, we turn attention to the six-component model, Equations (3) where both a primary tumor and a secondary organ are modeled. In Figures 4-6, we present bifurcation diagrams for the size of the tumor at the secondary site, 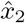, where it is assumed that the primary, well-vascularized (*I*_*LOMA*_, *J*_*LOMA*_ phenotype) tumor, 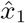, is eliminated 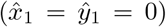. We therefore consider the equations for the second site,

**Fig. 4.**
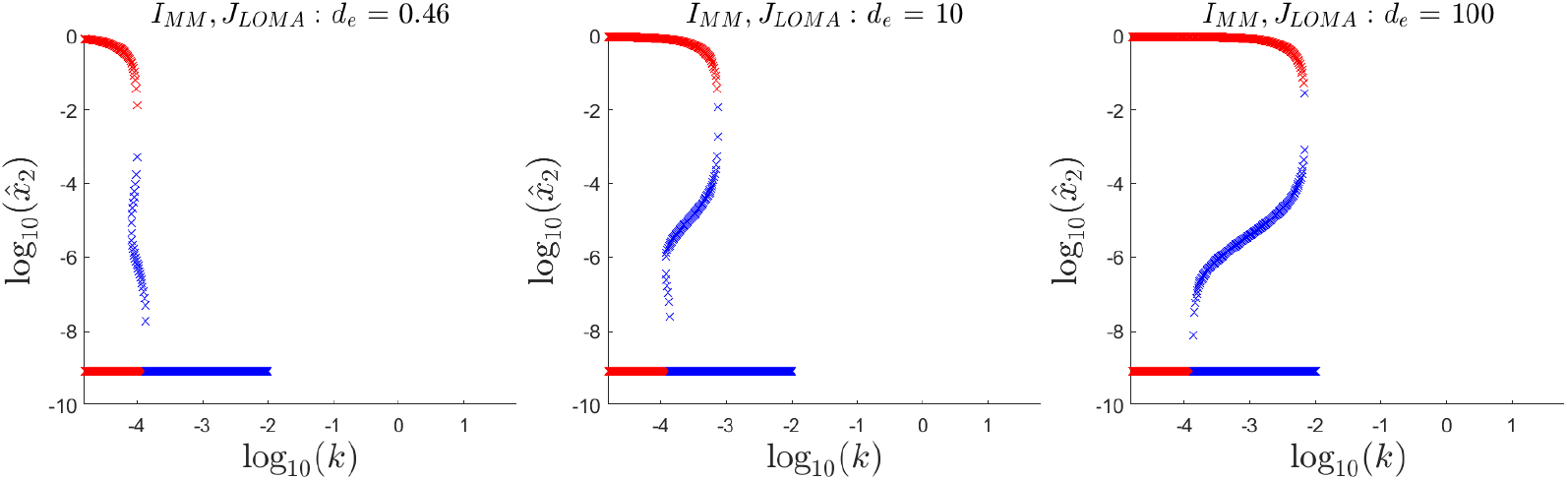
Bifurcation diagram for Equations (3) in the case that a primary tumor has been eliminated 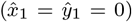. Stable equilibrium values of the secondary tumor 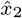 are colored blue and unstable values are colored red. All parameters are as shown in Table 1. We also follow [31, 33] in superimposing the tumor-free state on the logarithmic scaling by assigning it to a value of log_10_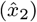less than all others. This branch, nonetheless, corresponds to 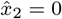.

**Fig. 5.**
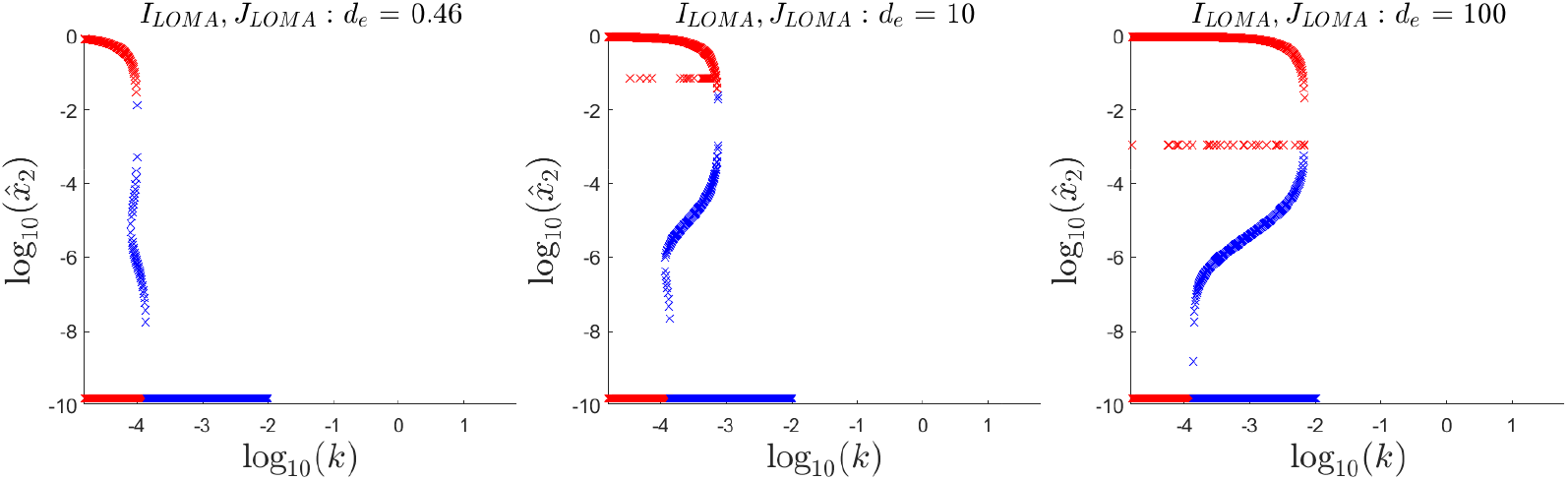
Bifurcation diagram for Equations (3) in the case that a primary tumor has been eliminated 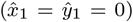. Stable equilibrium values of the secondary tumor 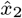 are colored blue and unstable values are colored red. All parameters are as shown in Table 1. We also follow [31, 33] in superimposing the tumor-free state on the logarithmic scaling by assigning it to a value of log_10_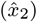 less than all others. This branch, nonetheless, corresponds to 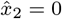.

**Fig. 6.**
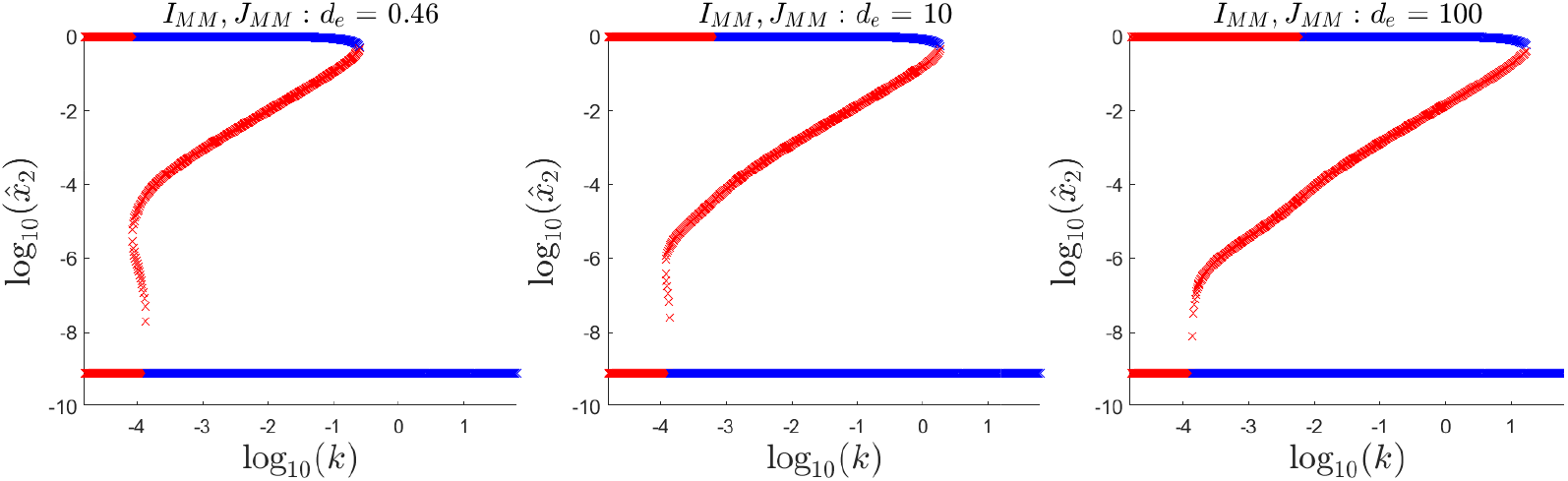
Bifurcation diagram for Equations (3) in the case that a primary tumor has been eliminated 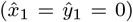. Stable equilibrium values of the secondary tumor 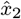 are colored blue and unstable values are colored red. As in the bifurcation diagram for the primary tumor alone (Figure 3), setting the infection rates *I* and killing rates *J* to Michaelis-Menten forms specializes the present model to the scenario of a poorly-vascularized tumor where the oncolytic virus and immune cells have only rate-limited access to tumor cells. All parameters are as shown in Table 1. We also follow [31, 33] in superimposing the tumor-free state on the logarithmic scaling by assigning it to a value of log_10_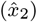 less than all others. This branch, nonetheless, corresponds to 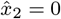.

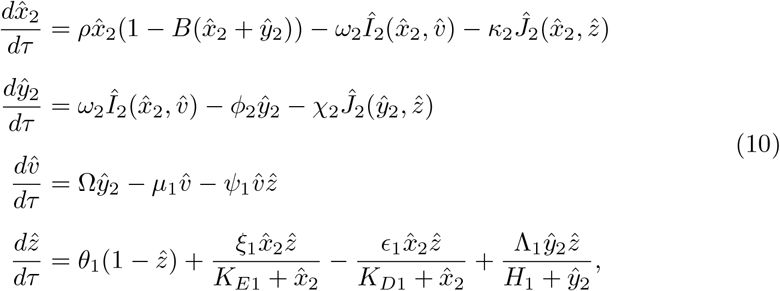

which are similar to Equations 9 up to parameter assignment. Setting 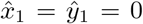 allows us to address one of our primary questions of interest — whether OVT can produce scenarios where a primary tumor is eliminated but metastasis to a secondary organ becomes favored — while also avoiding the intractable algebraic problem of systematically reducing Equations 3 to a small system of polynomial equations whose equilibria can be found with *vpasolve* without requiring prohibitive amounts of computing time.

Even before proceeding, some insight is possible. First, the similarity of Equations 9 and Equations 10 implies that neither 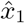nor 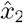 should have different equilibrium population sizes when the other site is assumed to be empty (or eliminated). However, it may be possible that some of the equilibrium tumor population values that are stable for 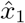in the single-site model, Equations 9, could be unstable for 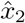in the two-site model, Equations 10 due to differences in the eigenvalues of the Jacobian — or vice-versa. The single-site model of Equations 9 leads to a Jacobian **J** ∈ ℝ^4×4^, expected to have 4 eigenvalues (counting multiplicity), while the two-site model of Equations 3 leads to a Jacobian **J** ∈ ℝ^6×6^, expected to have 6 eigenvalues (counting multiplicity). While this does not guarantee differences in stability between the two models, it suggests that differences *may* emerge.

We further note that assuming 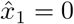 modifies the interpretation of the bifurcation diagrams. Strictly, the stability of an equilibrium value of 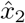, say, 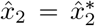 means that the equilibrium where *both* 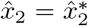 and 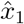= 0 are stable to perturbations. On the other hand, the instability of an equilibrium value of 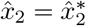 means that at *least* one of 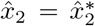 and 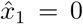 is unstable to perturbations. The bifurcation diagram alone cannot differentiate between the latter two cases. However, examination of the eigenvectors corresponding to the eigenvalue problem of Equation (8) reveals that *both* 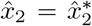 and 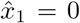 are unstable to perturbations. Hence the unstable branches of Figures 4-6 can be interpreted as corresponding to the inability of OVT to successfully destroy the primary tumor in a manner robust to noise.

We also point out that Equations (3) admits *both* 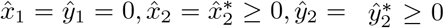 equilibria *and* 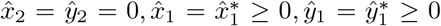 equilibria. This means that the equilibria represented in Figure 2 (where 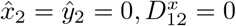) still exist in the six-dimensional phase space associated with Equations (3), and could ultimately attract trajectories originating from the unstable branches shown in Figures 4-6.

In Figure 4, we assume that the secondary tumor has functional responses *I*_*MM*_ and *J*_*LOMA*_. In Figure 5, we assume that the secondary tumor instead has functional responses *I*_*LOMA*_ and *J*_*LOMA*_. We comment on these two cases together because their bifurcation diagrams are nearly indistinguishable. Several observations are possible. First, the upper branch corresponding to a large secondary tumor even when the primary tumor has been destroyed 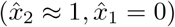 is unstable. As noted above, there are two possible interpretations: the equilibrium could be unstable because tumor 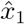 remains stable but tumor 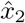 grows, *or* because both tumors 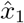and 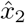 grow. In either case, the prognosis associated with the upper unstable branch is not positive. More certain is that stable, small-tumor equilibria exist for 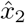 even when 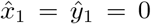. This provides an affirmative answer to our central question: whether it is possible for OVT to eliminate a primary tumor even while distant metastasis occurs to an *a priori* unknown location.

Figures 4 and 5 reveal unexpected dangers. The protocol suggested by, e.g., [31, 33, 34] entails OVT coupled with an initial phase of immune suppression (corresponding to increased *d*_*e*_ and possibly also decreased *k*) followed by immune boosting (corresponding to decreased *d*_*e*_ and possibly also increased *k*), possibly achieved through the use of checkpoint inhibitors [28, 31, 33, 34]. For moderate *k* in the range of normal physiology (*k ≈* 10^*™*4^; see Table 1), increasing *d*_*e*_ increases the interval of *k* where the unstable aggressive metastasis 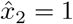 and unstable tumor-free 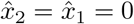 equilibria are directly opposed on the bifurcation diagram without the stable, smallmetastasis branch that occurs for smaller *d*_*e*_. This opposition of two unstable equilibria of high-dimensional systems suggests the possibility of oscillations or limit cycles. However, in the six-dimensional phase space of Equations (3), and given the presence of the other equilibria indicated by Figures 1-3, it is also natural to wonder whether these oscillations might plausibly allow the system to find an attracting but highly diseased state — especially in view of the fact that the six-component system evidently has more equilibria than the four-component system, some of which are attracting and correspond to well-developed metastases. Hence Figures 4 and 5 suggest two important possible failure modes of OVT coupled with immune modulation: (i) producing oscillations that ultimately lead to the system to settle in a state where both the primary and secondary tumors are able to grow without bound; or (ii) that the restoration of normal immune function produces trajectories settling in a state where the primary tumor is cleared but there exist secondary metastases at subclinical (or nearly subclinical) concentrations 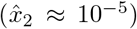 which may lead to fully-developed tumors later.

In Figure 6, we consider metastasis to a poorly vascularized secondary organ. As with the case of a primary tumor with metastasis neglected, the bifurcation diagram of phenotype *I*_*MM*_, *J*_*MM*_ is exceptional, and several aspects are particularly striking. First, there is again the possibility that undesirable behavior occurs when immune suppression is reversed (i.e., decreasing *d*_*e*_): in contrast with the *I*_*MM*_, *J*_*LOMA*_ and *I*_*LOMA*_, *J*_*LOMA*_ phenotypes for the secondary tumor for which the large-metastasis branch is always unstable, there are newly parts of the upper, large-metastasis branch that are stable and lie above an *unstable* small-metastasis branch for a wide range of *k*. This means that it is no longer just a possibility that the primary tumor is eliminated while metastasis to a secondary organ is promoted — it is the attracting state for a comparatively large part of the phase space. The possibility that decreasing *d*_*e*_ induces limit cycles for *k ≈* 10^*™*4^ persists.

Beyond the present mathematical idealization, a reality that cannot possibly be captured by bifurcation diagrams is that real human physiology is in a state of dynamic equilibrium. One does not expect that all simulation parameters are truly constants. Moreover, tumor vascularization is known to be dynamic, and tumors can change phenotypes. What appears to be a single tumor mass is often, in reality, an aggregate of various tumor phenotypes competing for resources. Accordingly, we emphasize that the changes in the bifurcation diagrams are of qualitative interest even if we have assumed for the simplicity of modeling that the parameters of Table 1 are knowable *a priori* and fixed.

The modern oncological toolkit includes vascular disrupting agents (VDAs), which, in the present modeling framework, would be expected to rapidly convert an existing tumor to an *I*_*MM*_, *J*_*MM*_ phenotype, greatly modifying its response to OVT if administered simultaneously. To this point, Figures 4-6 suggest that there exist regions of parameter space where the sudden transformation of a tumor from well-vascularized *I*_*MM*_, *J*_*LOMA*_ or *I*_*LOMA*_, *J*_*LOMA*_ type to poorly-vascularized *I*_*MM*_, *J*_*MM*_ type could lead to unfavorable patient outcomes, with trajectories becoming trapped in regions where aggressive metastasis is the only attracting state. In Section 5, we briefly investigate this hypothesis.

## 5 Results: simulation studies

The bifurcation analysis of Sections 4.1-4.2 indicates that, in general, when tumors are sufficiently small or the immune killing rate is sufficiently large, a tumor is eliminated by the immune system. On the other hand, if the immune killing rate is sufficiently small, a variety of dangerous phenomena can occur, including uncontrolled growth of a primary tumor, metastasis to a secondary organ, and possibly limit cycles in phase space corresponding to primary and secondary tumors whose size oscillates about some average over time.

In this Section, we consider the liminal scenario where *k* = 10^*™*4^ and *d*_*e*_ = 0.46, there is no initial secondary tumor 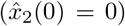 and we run a large suite of simulations sweeping over (i) initial primary tumor sizes 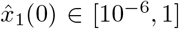 when treatment begins and (ii) the dosage of oncolytic virus administered at the time 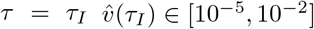 We assign the initial dimensionless immune cell concentration as 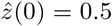, which lies in between the two extremes 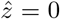, representing total immune failure, and 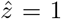, which occurs in the tumor-free state. Initial infected cell populations are assigned as 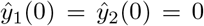. At *τ*_*I*_, the dimensionless virion count 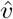 is assigned to the prescribed dose. All other parameters, unless otherwise specified, are the same as in Table 1. Although it is impossible to consider every variation on a clinical protocol combining OVT, immune modulation, and VDAs, we simulate here three hypothetical, highly idealized approaches to treatment:

- **Protocol I: immune suppression first, then OVT**. Following the two-phase immune modulation protocol suggested by, e.g., [31, 33, 34], immune suppressants are administered at dimensionless time *τ* = 0, increasing *d*_*e*_ to 100. Then, at *τ* = 2, we set 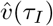 to the specified dose. At *τ* = 4, immune suppression is reversed and *d*_*e*_ is set back to 0.46.
- **Protocol II: concurrent immune suppression and OVT**. Here, we instead apply OVT and immune suppressants are administered at dimensionless time *τ*_*I*_ = 0. At this time, we set 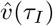 ) to the specified dose and *d*_*e*_ to 100, representing immune suppression. At *τ* = 4, immune suppression is reversed and *d*_*e*_ is set back to 0.46.
- **Protocol III: OVT first, then VDA**. At *τ* = 0, we set 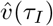 to the speci-fied dose to debulk the primary tumor. At *τ* = 2, a VDA is administered. We assume that the VDA has the immediate effect of converting both the primary and secondary tumors to phenotype *I*_*MM*_, *J*_*MM*_, regardless of their initial phenotype, representing a poorly-vascularized tumor. Although somewhat unrealistic, in part because VDAs are not generally used as front-line treatments, this will allow an assessment of whether it is the case, as suggested by the bifurcation diagrams, that devascularization might unintentionally stabilize a tumor against treatment by OVT.

For the sake of comparison, we also establish two baseline treatments:

- **Baseline I: No treatment**. In this case, no intervention of any kind is made.
- **Baseline II: OVT alone**. In this case, we apply OVT at *τ* = 0 according to the specified dose, 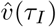, but there is no immune suppression.

In all cases, the simulation is terminated at *τ* = 20, corresponding to around 65 days given *r* = 0.346 (see Table 1). When immune modulation is simulated, the delay of Δ*τ* = 2 (around seven days for *r* = 0.346) between the administration of immune suppression and OVT is motivated by recent clinical studies [77–79] where a comparable protocol was used. The Δ*τ* = 20 duration of simulations is intended to reasonably coincide with ongoing clinical studies where treatment cycles might last a few weeks; see [16, 25, 32, 80] and other references therein. We aim to bypass the questions of optimal scheduling and long-term prognosis in favor of modeling the state of the tumor following *a single treatment cycle*.

Hence, in Figures 7-20, we present heat maps of the final populations of the primary and secondary tumors, 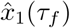 and 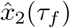 respectively, plotted against the administered virion population and the initial primary tumor size. For some of these simulation studies, we also produce a small number of representative time series plots. In what follows, we first present all figures, and then comment on all of them at once.

### Comments on numerical simulations

Simulations of Equations (3) are implemented using Matlab’s *ode15s*. These simulations require careful attention to two details that were comparatively simple to handle in Sections 4.1-4.2.

First, *simulation outputs for* 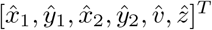 *must never be negative*. This is easily handled within Matlab’s *ode15s* by turning on the “NonNegative” option.

Second, *populations corresponding to less than a single virion or cell are nonphysical*. While simulations of Equations (3) are continuous and can produce dimensionless quantities corresponding to, e.g., 10^*™*2^ of a cell, such populations are nonphysical — or, at least, their physical meaning is difficult to interpret. It is equally nonphysical to suppose that an entire tumor can regrow from a surviving population corresponding to less than one cell. This is sometimes referred to as the “yocto-cell” problem [81]. To enforce physically-valid outputs, we apply thresholding: any model output for the dimensionless quantities 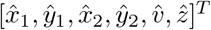 corresponding to less than 1 unit has the autonomous growth terms in its evolution equation immediately set to zero, as are metastatic seeding terms 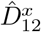 when 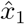 falls below threshold. See the nondimensional-ization scheme of Section 3. For example, if simulations of Equations (3) ever produce 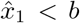, we set the logistic growth term 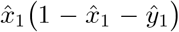 to zero, preventing regeneration. Its shedding via the term 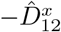, and the corresponding source term 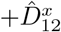 in the evolution equation for 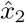, are also set to zero. This ensures that populations that have gone extinct stay extinct, with two possible exceptions: (i) immune cells 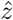 are constantly recruited (see Equations 2), and (ii) a secondary tumor 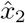 of size zero could be resuscitated by seeding from an extant primary tumor 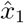.

### 5.1 Baseline I: No treatment

**Fig. 7.**
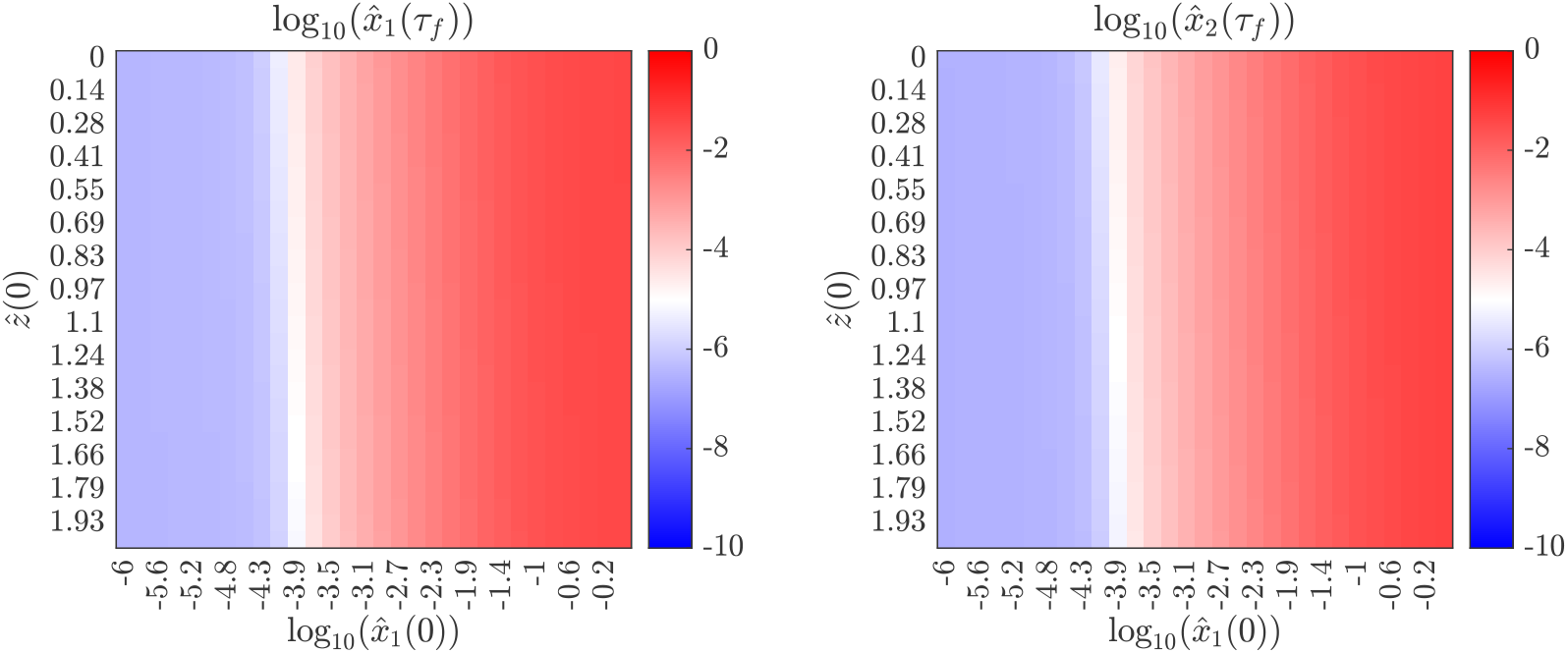
Simulation results at *τ* = 20 with an initial tumor of dimensionless size 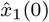. To establish a baseline, no treatment is given. The primary tumor is assumed to have *I*_*LOMA*_, *J*_*LOMA*_ phenotype and metastases are assumed to have phenotype *I*_*LOMA*_, *J*_*LOMA*_. **Left:** The value of 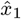, the primary tumor size, at *τ* = 20, the end of the simulation. **Right:** The value of 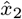, the secondary tumor, at *τ* = 20, the end of the simulation.

**Fig. 8.**
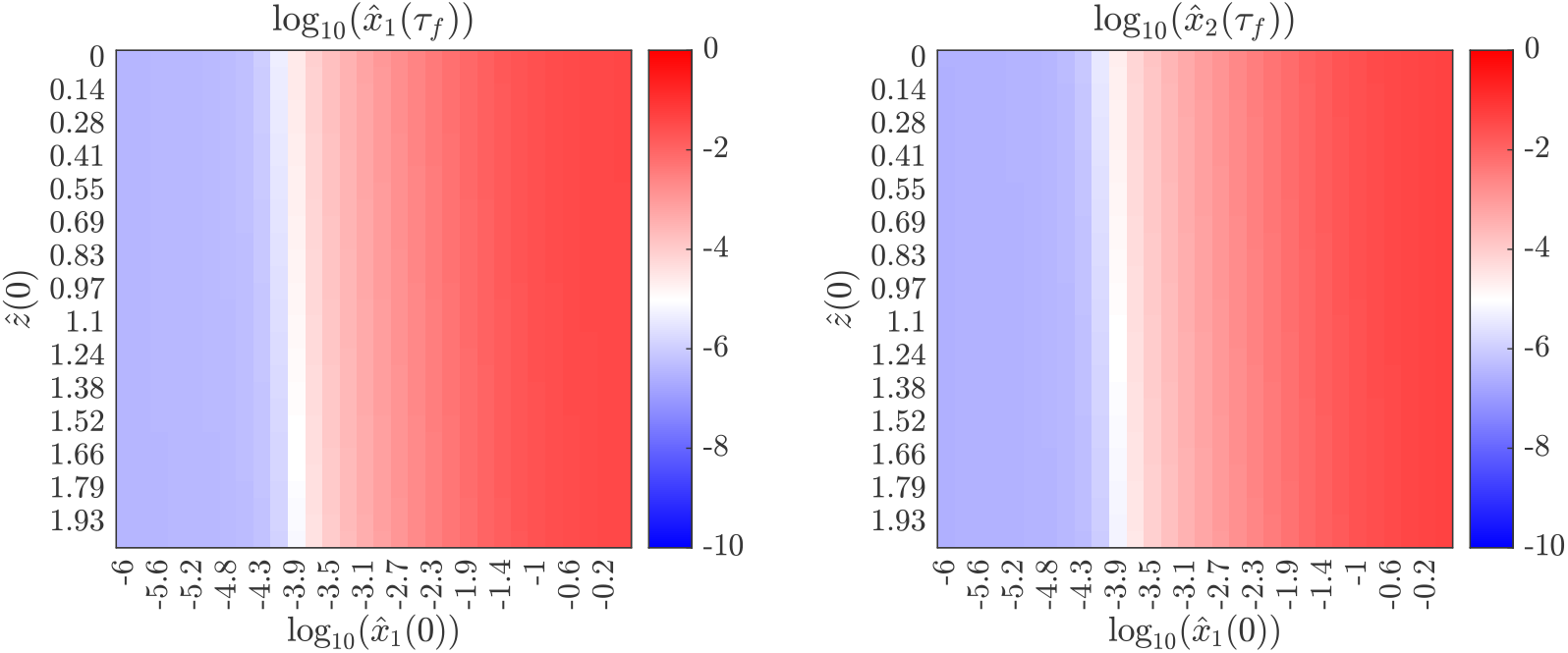
Simulation results at *τ* = 20 with an initial tumor of dimensionless size 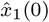. To establish a baseline, no treatment is given. The primary tumor is assumed to have *I*_*LOMA*_, *J*_*LOMA*_ phenotype and metastases are assumed to have phenotype *I*_*MM*_, *J*_*LOMA*_. **Left:** The value of 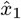, the primary tumor size, at *τ* = 20, the end of the simulation. **Right:** The value of 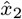, the secondary tumor, at *τ* = 20, the end of the simulation.

**Fig. 9.**
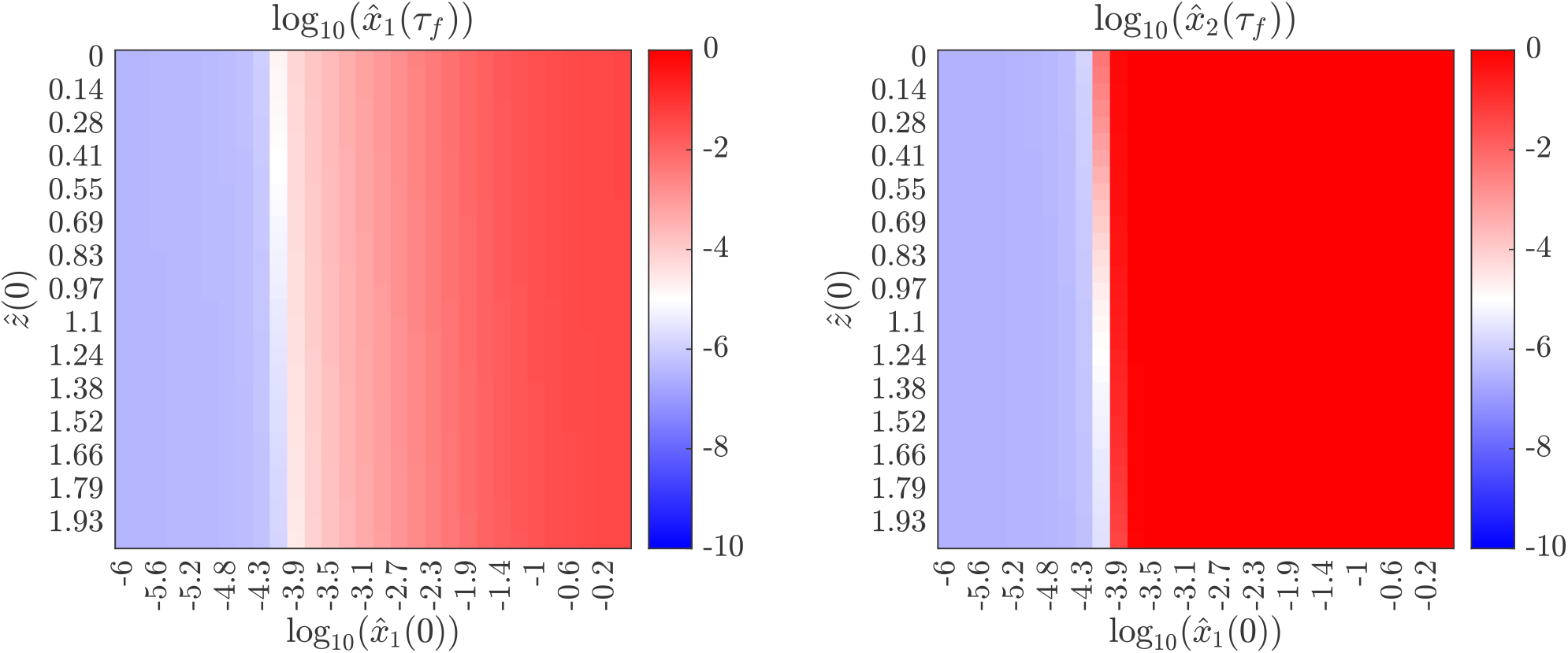
Simulation results at *τ* = 20 with an initial tumor of dimensionless size 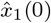. To establish a baseline, no treatment is given. The primary tumor is assumed to have *I*_*LOMA*_, *J*_*LOMA*_ phenotype and metastases are assumed to have phenotype *I*_*MM*_, *J*_*MM*_ . **Left:** The value of 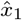, the primary tumor size, at *τ* = 20, the end of the simulation. **Right:** The value of 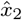. the secondary tumor, at *τ* = 20, the end of the simulation.

### 5.2 Baseline II: OVT without immune modulation

**Fig. 10.**
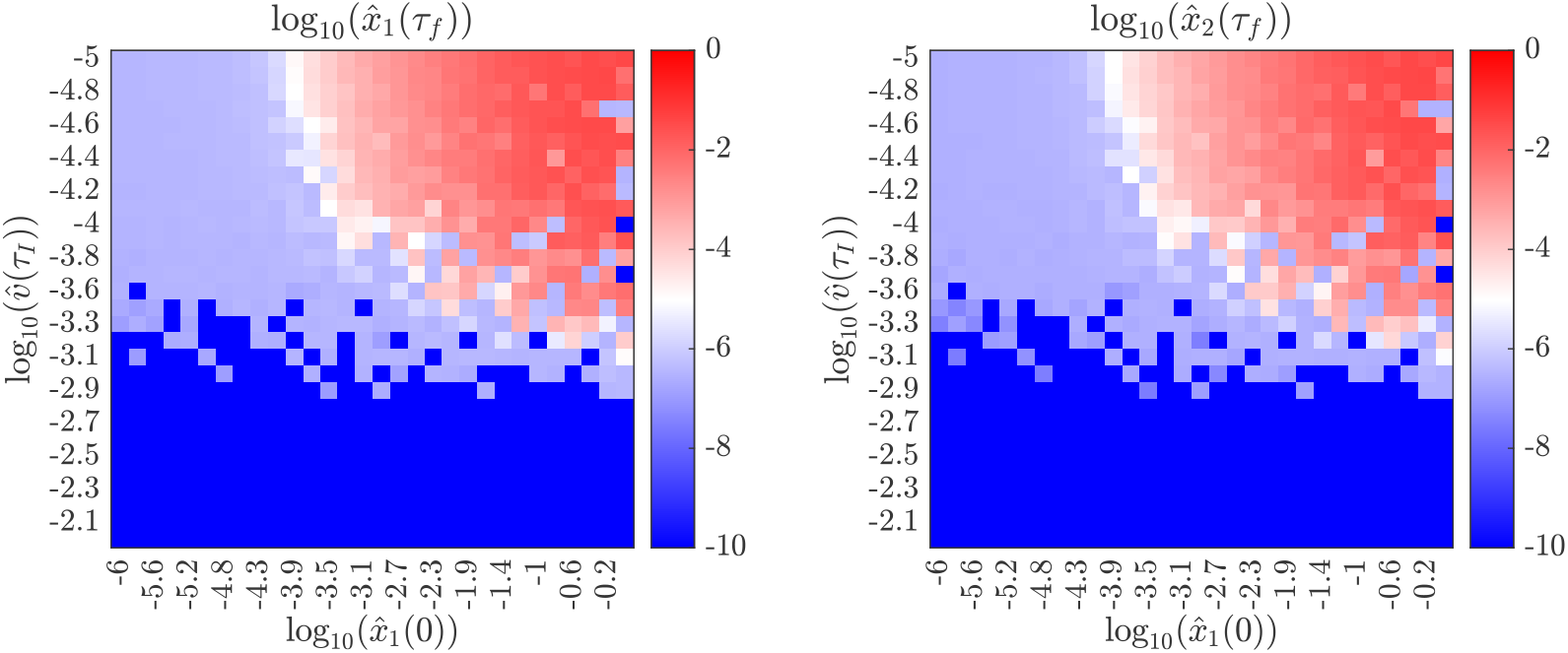
Simulation results at *τ* = 20 following injection of 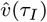 dimensionless virions at *τ*_*I*_ = 0, and an initial tumor of dimensionless size 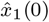. The primary tumor is assumed to have *I*_*LOMA*_, *J*_*LOMA*_ phenotype and metastases are assumed to have phenotype *I*_*LOMA*_, *J*_*LOMA*_. **Left:** The value of 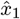, the primary tumor size, at *τ* = 20, the end of the simulation. **Right:** The value of 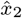, the secondary tumor, at *τ* = 20, the end of the simulation.

**Fig. 11.**
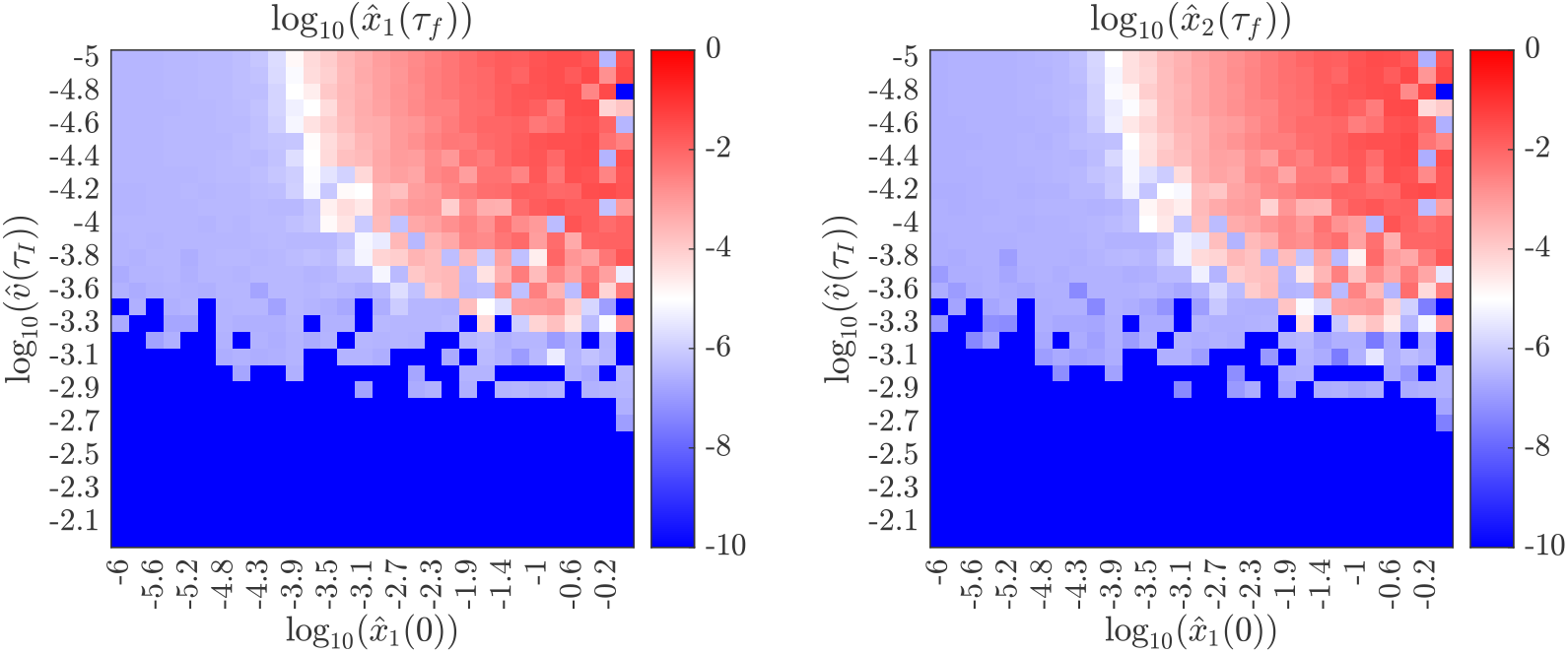
Simulation results at *τ* = 20 following injection of 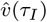 dimensionless virions at *τ*_*I*_ = 0, and an initial tumor of dimensionless size 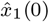. The primary tumor is assumed to have *I*_*LOMA*_, *J*_*LOMA*_ phenotype and metastases are assumed to have phenotype *I*_*MM*_, *J*_*LOMA*_. **Left:** The value of 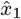, the primary tumor size, at *τ* = 20, the end of the simulation. **Right:** The value of 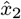, the secondary tumor, at *τ* = 20, the end of the simulation.

**Fig. 12.**
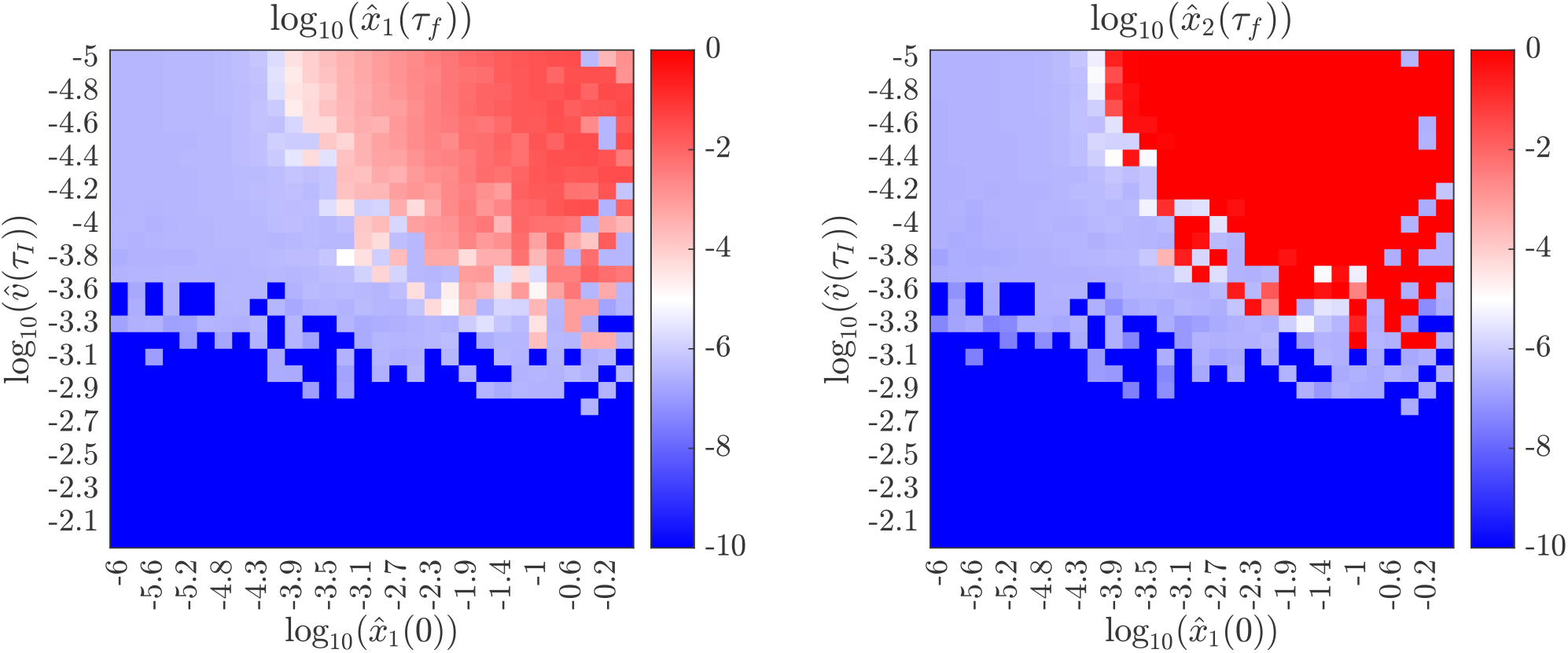
Simulation results at *τ* = 20 following injection of 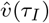 dimensionless virions at *τ*_*I*_ = 0, and an initial tumor of dimensionless size 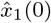. The primary tumor is assumed to have *I*_*LOMA*_, *J*_*LOMA*_ phenotype and metastases are assumed to have phenotype *I*_*MM*_, *J*_*MM*_ . **Left:** The value of 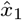., the primary tumor size, at *τ* = 20, the end of the simulation. **Right:** The value of 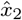, the secondary tumor, at *τ* = 20, the end of the simulation.

### 5.3 Protocol I: immune suppression followed by OVT

**Fig. 13.**
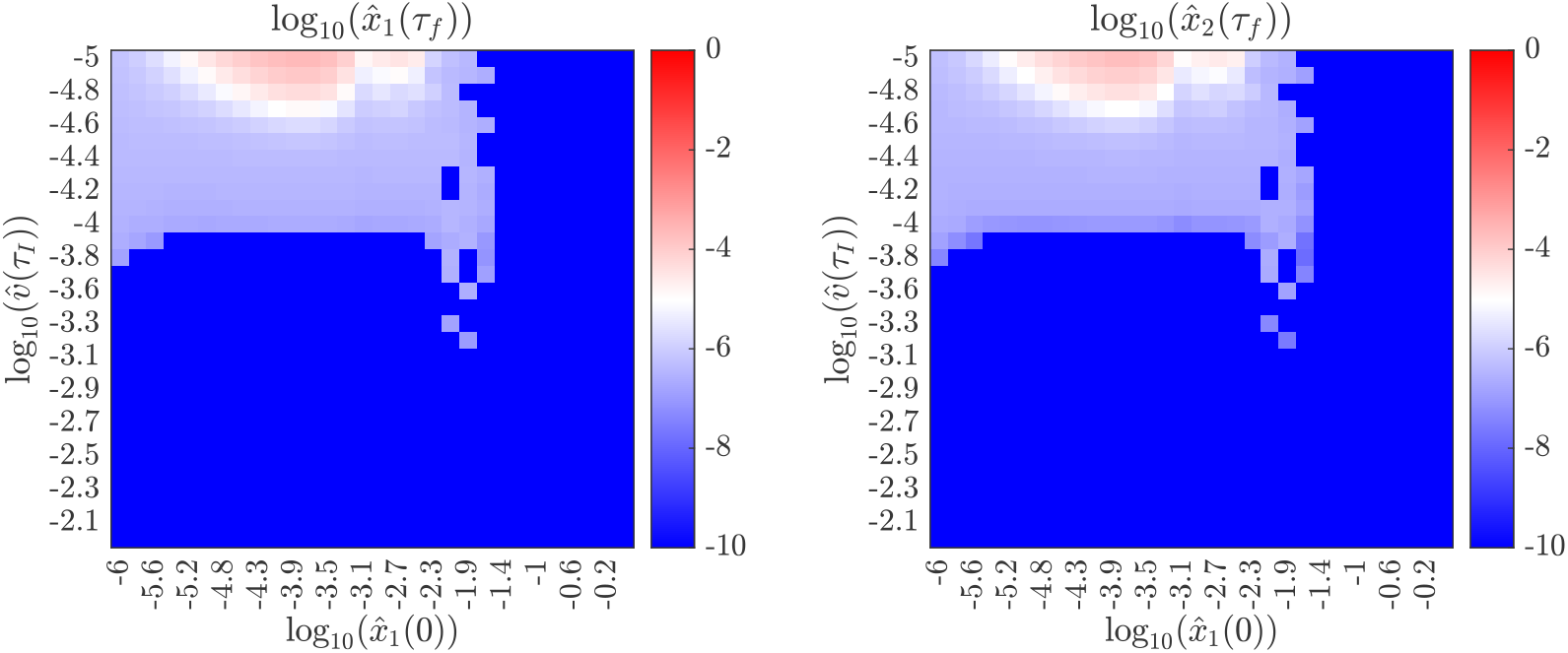
Simulation results at *τ* = 20 after setting *d*_*e*_ = 100 at *τ* = 0, injection of 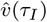 dimensionless virions at *τ*_*I*_ = 2, and an initial tumor of dimensionless size 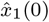. At *τ* = 4, we set *d*_*e*_ = 0.46, modeling the restoration of normal immune function. The primary tumor is assumed to have *I*_*LOMA*_, *J*_*LOMA*_ phenotype and metastases are assumed to have phenotype *I*_*LOMA*_, *J*_*LOMA*_. **Left:** The value of 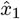, the primary tumor size, at *τ* = 20, the end of the simulation. **Right:** The value of 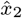, the secondary tumor, at *τ* = 20, the end of the simulation. This figure coincides with Protocol I.

**Fig. 14.**
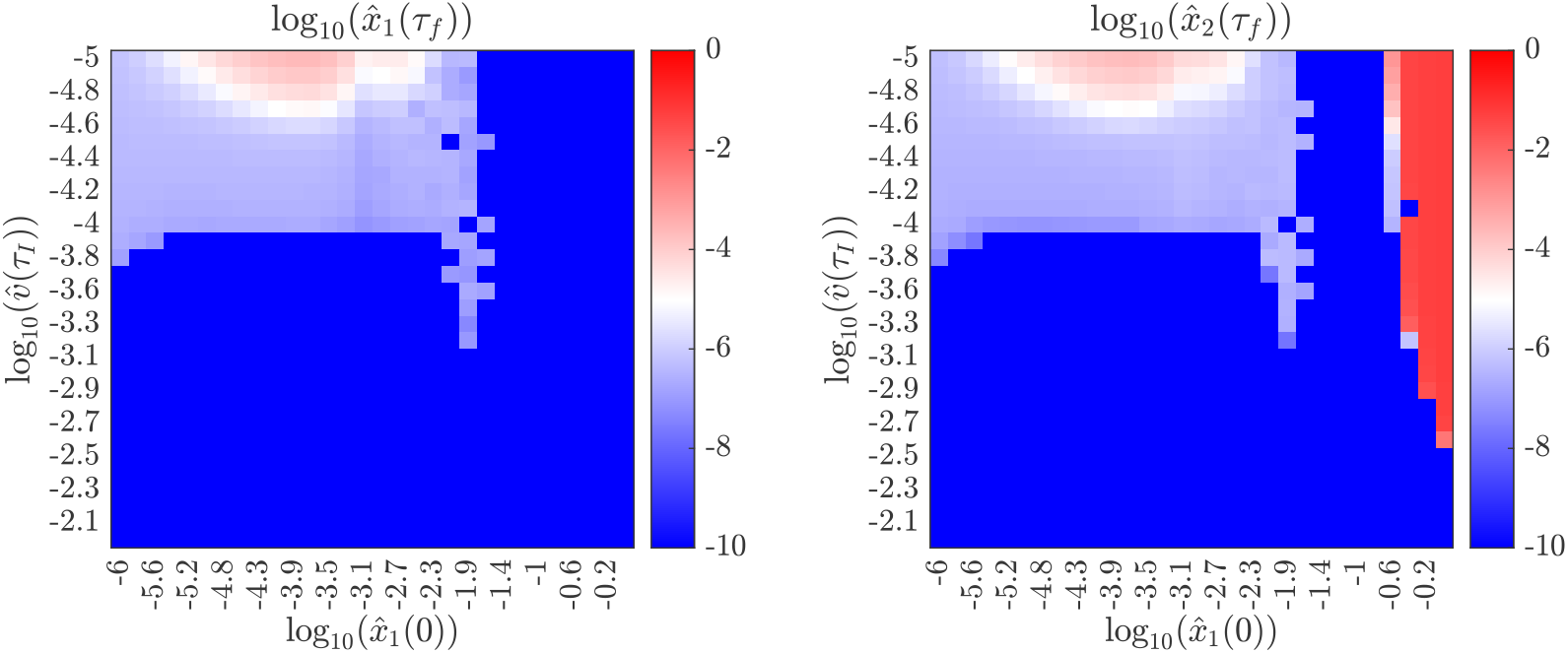
Simulation results at *τ* = 20 after setting *d*_*e*_ = 100 at *τ* = 0, injection of 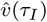 dimensionless virions at *τ*_*I*_ = 2, and an initial tumor of dimensionless size 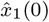. At *τ* = 4, we set *d*_*e*_ = 0.46, modeling the restoration of normal immune function. The primary tumor is assumed to have *I*_*LOMA*_, *J*_*LOMA*_ phenotype and metastases are assumed to have phenotype *I*_*MM*_, *J*_*LOMA*_. **Left:** The value of 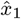, the primary tumor size, at *τ* = 20, the end of the simulation. **Right:** The value of 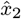, the secondary tumor, at *τ* = 20, the end of the simulation. This figure coincides with Protocol I.

**Fig. 15.**
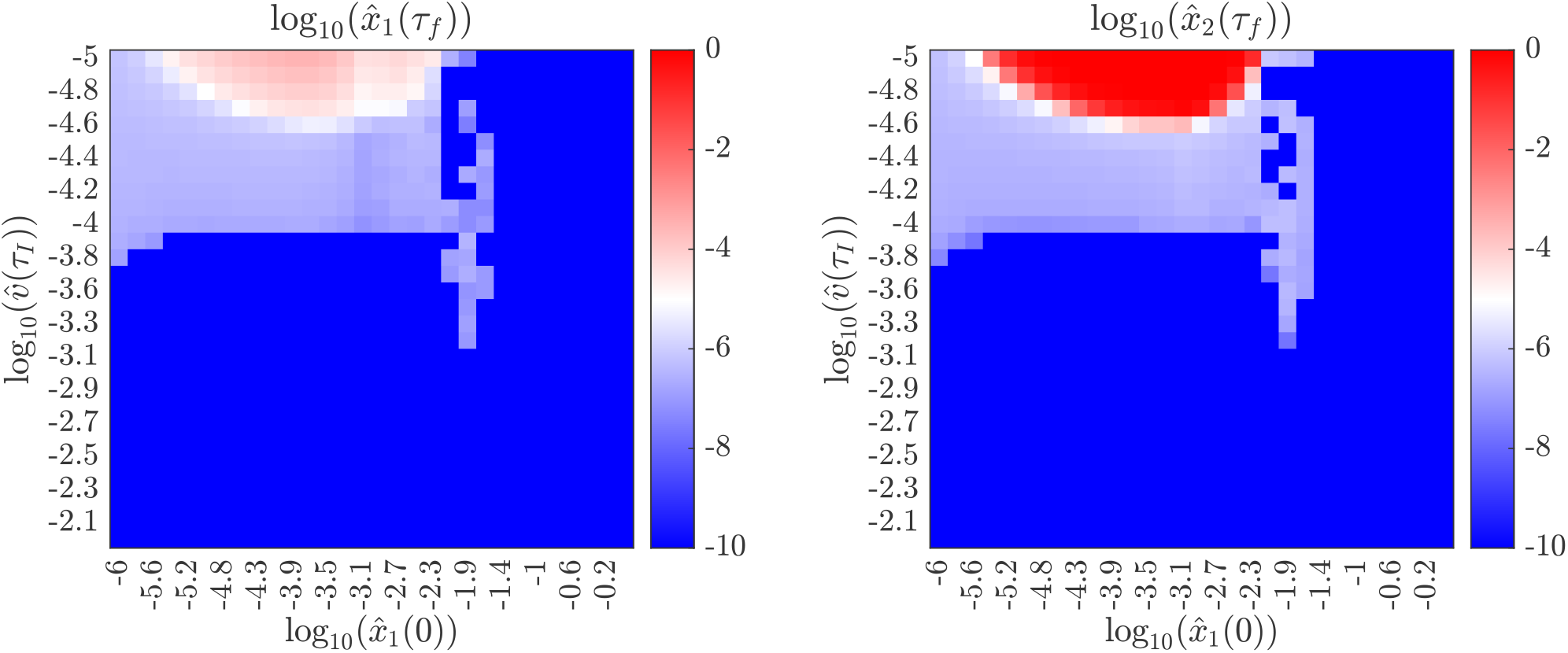
Simulation results at *τ* = 20 after setting *d*_*e*_ = 100 at *τ* = 0, injection of 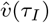 dimensionless virions at *τ*_*I*_ = 2, and an initial tumor of dimensionless size 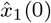. At *τ* = 4, we set *d*_*e*_ = 0.46, modeling the restoration of normal immune function. The primary tumor is assumed to have *I*_*LOMA*_, *J*_*LOMA*_ phenotype and metastases are assumed to have phenotype *I*_*MM*_, *J*_*MM*_ . **Left:** The value of 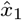, the primary tumor size, at *τ* = 20, the end of the simulation. **Right:** The value of 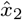, the secondary tumor, at *τ* = 20, the end of the simulation. This figure coincides with Protocol I.

### 5.4 Protocol II: simultaneous immune suppression and OVT

**Fig. 16.**
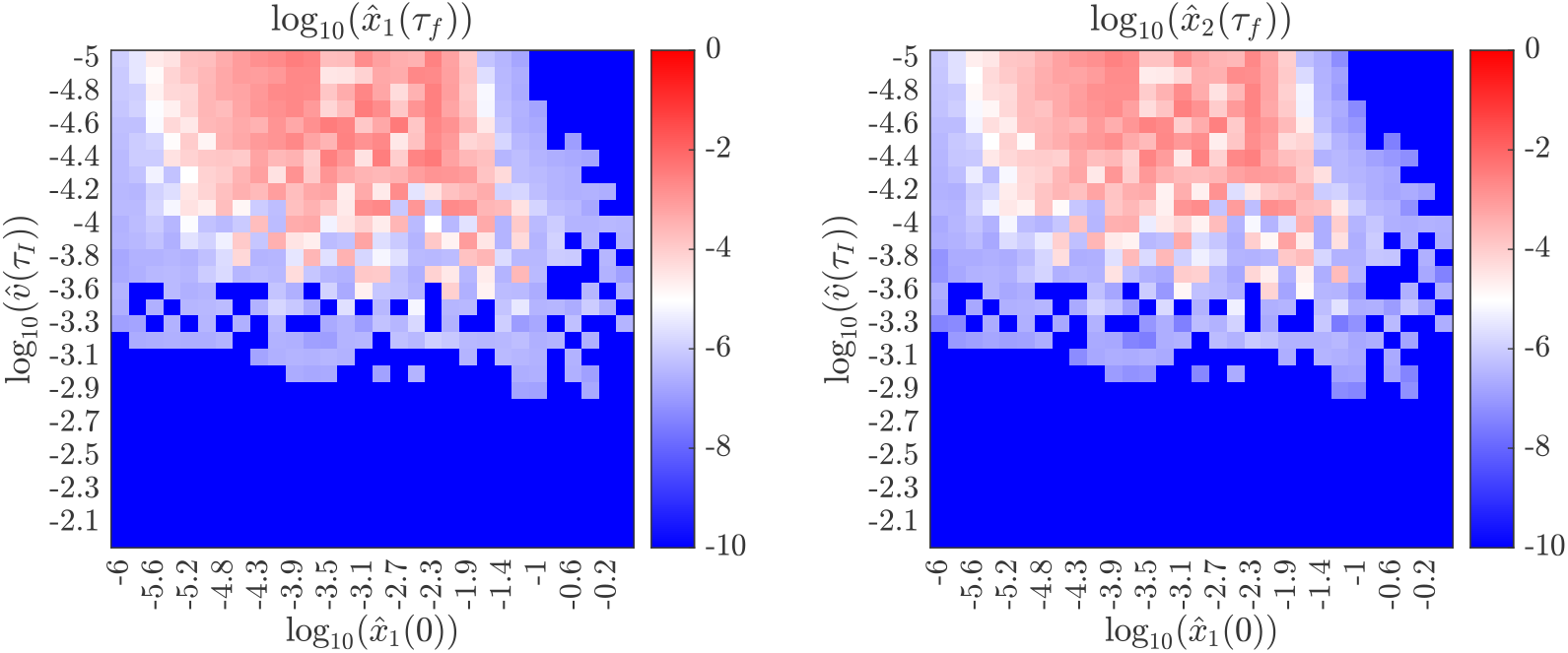
Simulation results at *τ* = 20 after setting *d*_*e*_ = 100 and injection of 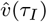 dimensionless virions, both at *τ*_*I*_ = 0, and an initial tumor of dimensionless size 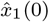. At *τ* = 4, we set *d*_*e*_ = 0.46, modeling the restoration of normal immune function. The primary tumor is assumed to have *I*_*LOMA*_, *J*_*LOMA*_ phenotype and metastases are assumed to have phenotype *I*_*LOMA*_, *J*_*LOMA*_. **Left:** The value of 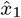, the primary tumor size, at *τ* = 20, the end of the simulation. **Right:** The value of 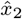, the secondary tumor, at *τ* = 20, the end of the simulation. This figure coincides with Protocol II.

**Fig. 17.**
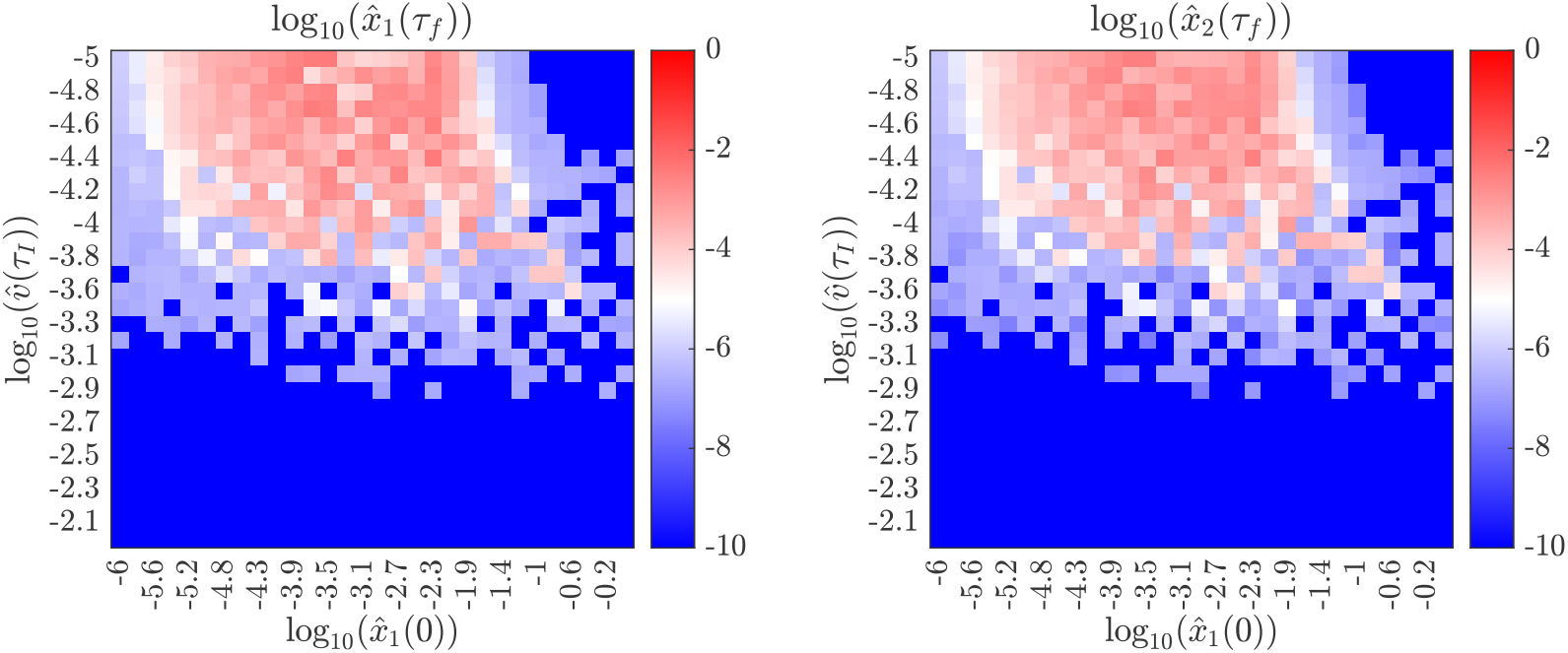
Simulation results at *τ* = 20 after setting *d*_*e*_ = 100 and injection 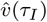 dimensionless virions, both at *τ*_*I*_ = 0, and an initial tumor of dimensionless size 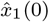.At *τ* = 4, we set *d*_*e*_ = 0.46, modeling the restoration of normal immune function. The primary tumor is assumed to have *I*_*LOMA*_, *J*_*LOMA*_ phenotype and metastases are assumed to have phenotype *I*_*MM*_, *J*_*LOMA*_. **Left:** The value of 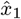, the primary tumor size, at *τ* = 20, the end of the simulation. **Right:** The value of 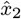, the secondary tumor, at *τ* = 20, the end of the simulation. This figure coincides with Protocol II.

**Fig. 18.**
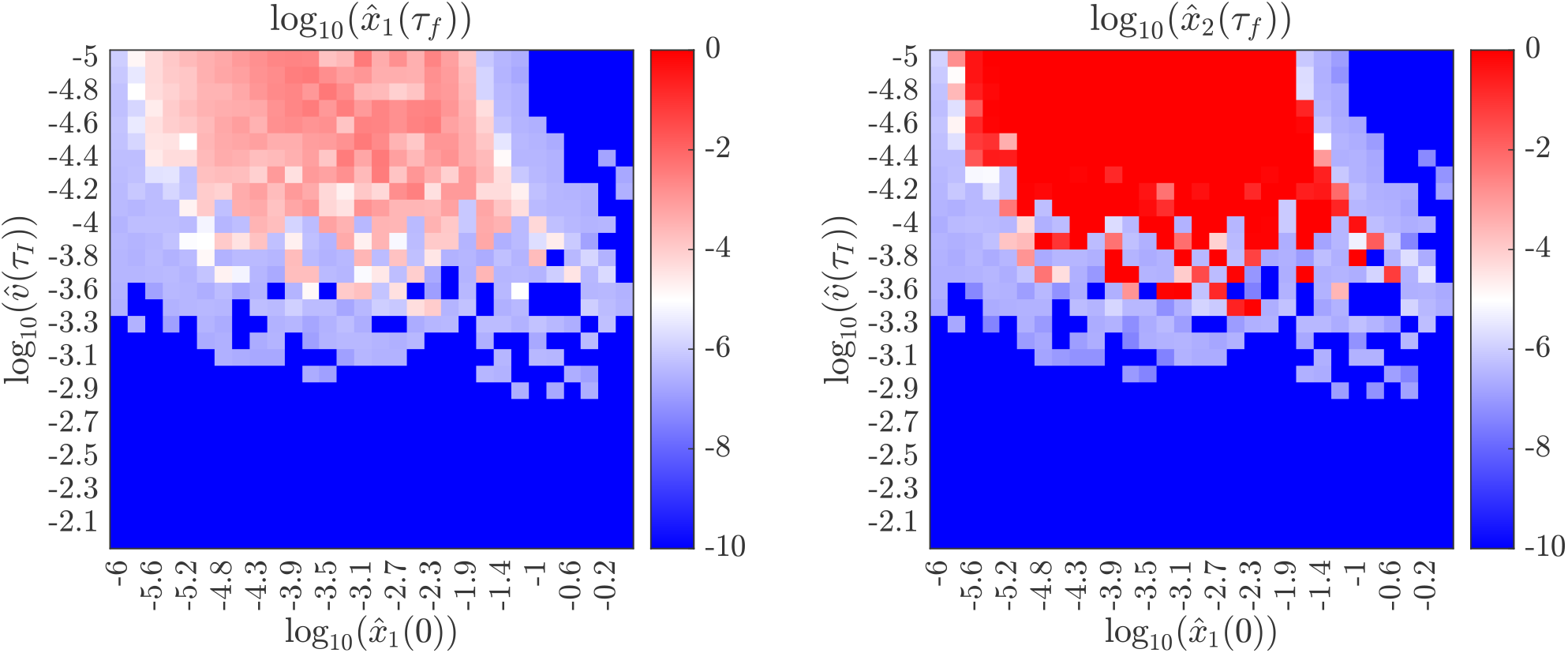
Simulation results at *τ* = 20 after setting *d*_*e*_ = 100 and injection of 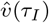 dimensionless virions, both at *τ*_*I*_ = 0, and an initial tumor of dimensionless size 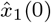 At *τ* = 4, we set *d*_*e*_ = 0.46, modeling the restoration of normal immune function. The primary tumor is assumed to have *I*_*LOMA*_, *J*_*LOMA*_ phenotype and metastases are assumed to have phenotype *I*_*MM*_, *J*_*MM*_ . **Left:** The value of 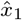, the primary tumor size, at *τ* = 20, the end of the simulation. **Right:** The value of 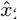, the secondary tumor, at *τ* = 20, the end of the simulation. This figure coincides with Protocol II.

### 5.5 Protocol III: OVT and vascular disrupting agents

**Fig. 19.**
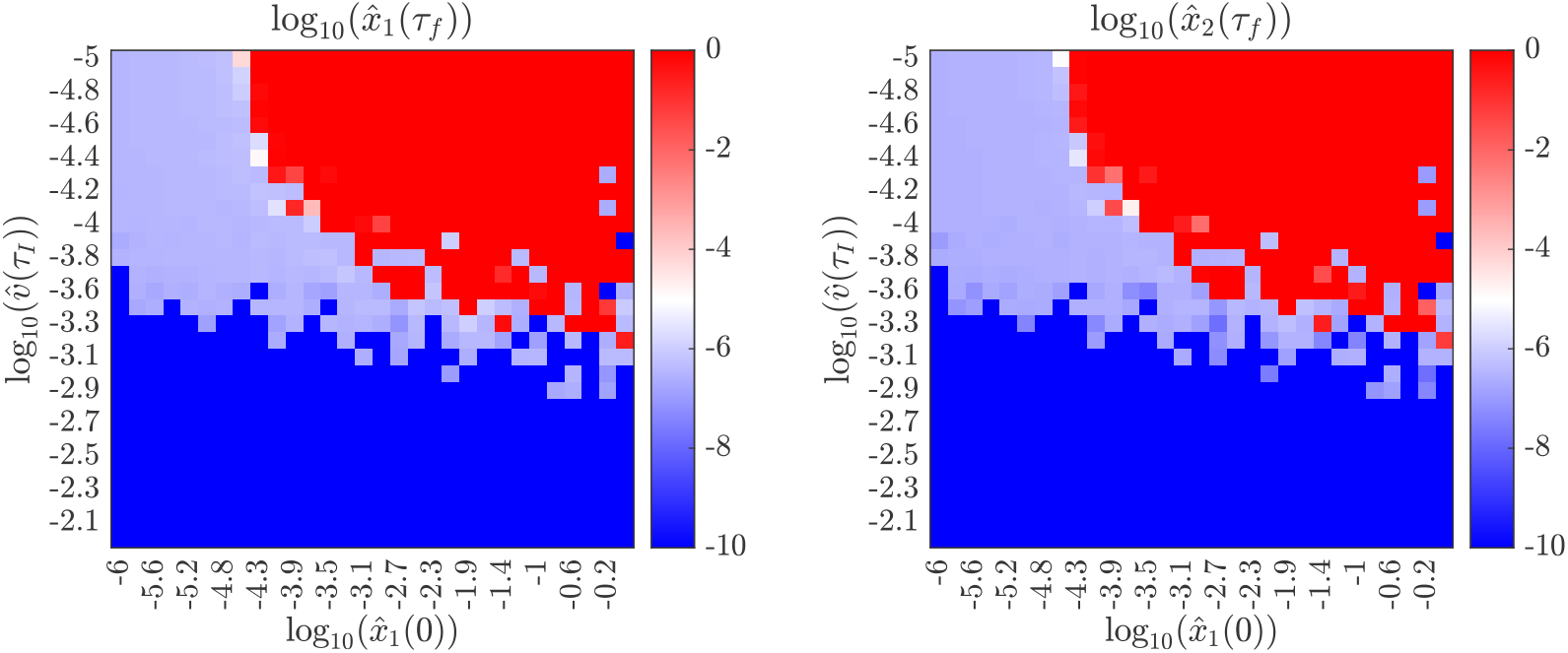
Simulation results at *τ* = 20 after injection of 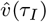dimensionless virions at *τ*_*I*_ = 0, and an initial tumor of dimensionless size 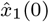. At *τ* = 2, we apply a VDA, modeled as the immediate and irreversible conversion of both the primary tumor and secondary tumor to *I*_*MM*_, *J*_*MM*_ phenotypes. The primary tumor is assumed to have initial phenotype *I*_*LOMA*_, *J*_*LOMA*_ and metastases are assumed to have initial phenotype *I*_*LOMA*_, *J*_*LOMA*_. **Left:** The value of 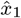, the primary tumor size, at *τ* = 20, the end of the simulation. **Right:** The value of 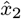, the secondary tumor, at *τ* = 20, the end of the simulation. This figure coincides with Protocol III.

**Fig. 20.**
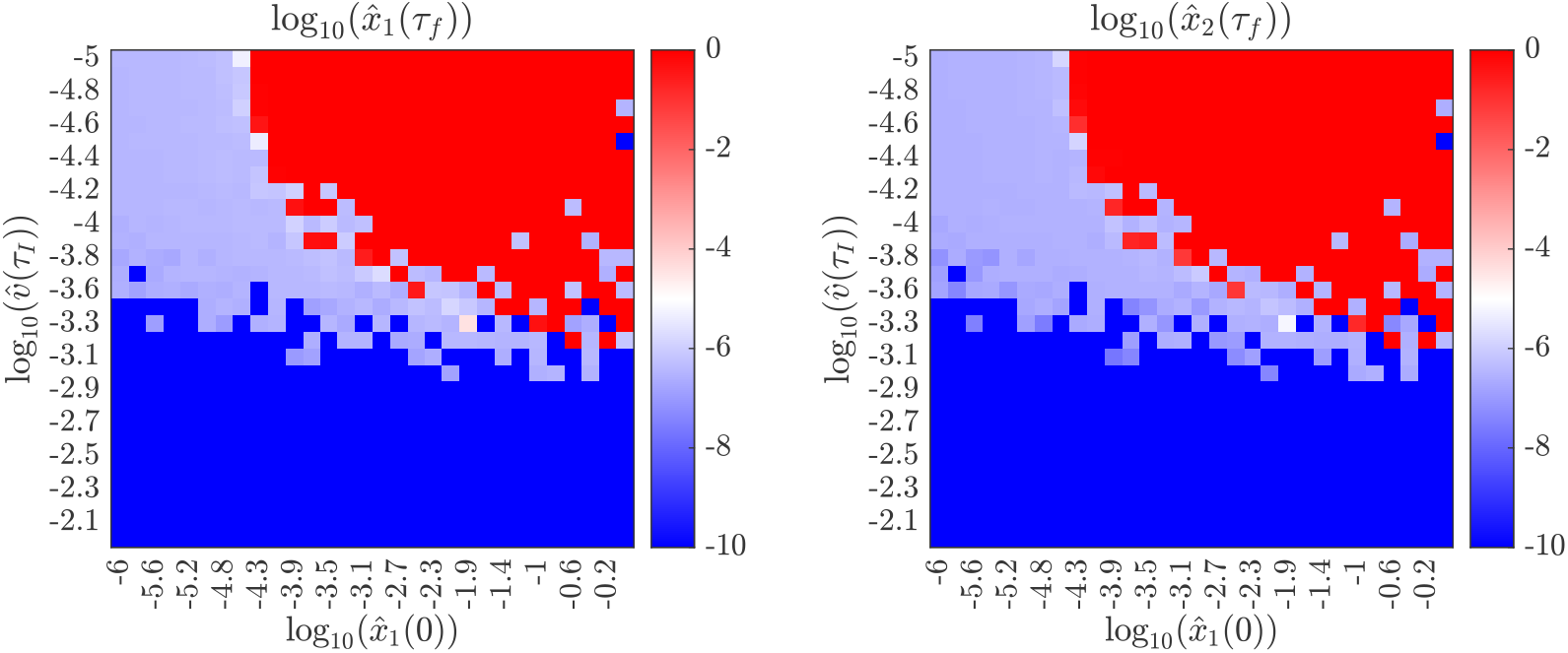
Simulation results at *τ* = 20 after injection of 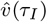 dimensionless virions at *τ*_*I*_ = 0, and an initial tumor of dimensionless size 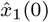. At *τ* = 2, we apply a VDA, modeled as the immediate and irreversible conversion of both the primary tumor and secondary tumor to *I*_*MM*_, *J*_*MM*_ phenotypes. The primary tumor is assumed to have initial phenotype *I*_*LOMA*_, *J*_*LOMA*_ and metastases are assumed to have initial phenotype *I*_*MM*_, *J*_*LOMA*_. **Left:** The value of 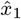, the primary tumor size, at *τ* = 20, the end of the simulation. **Right:** The value of 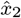, the secondary tumor, at *τ* = 20, the end of the simulation. This figure coincides with Protocol III.

**Fig. 21.**
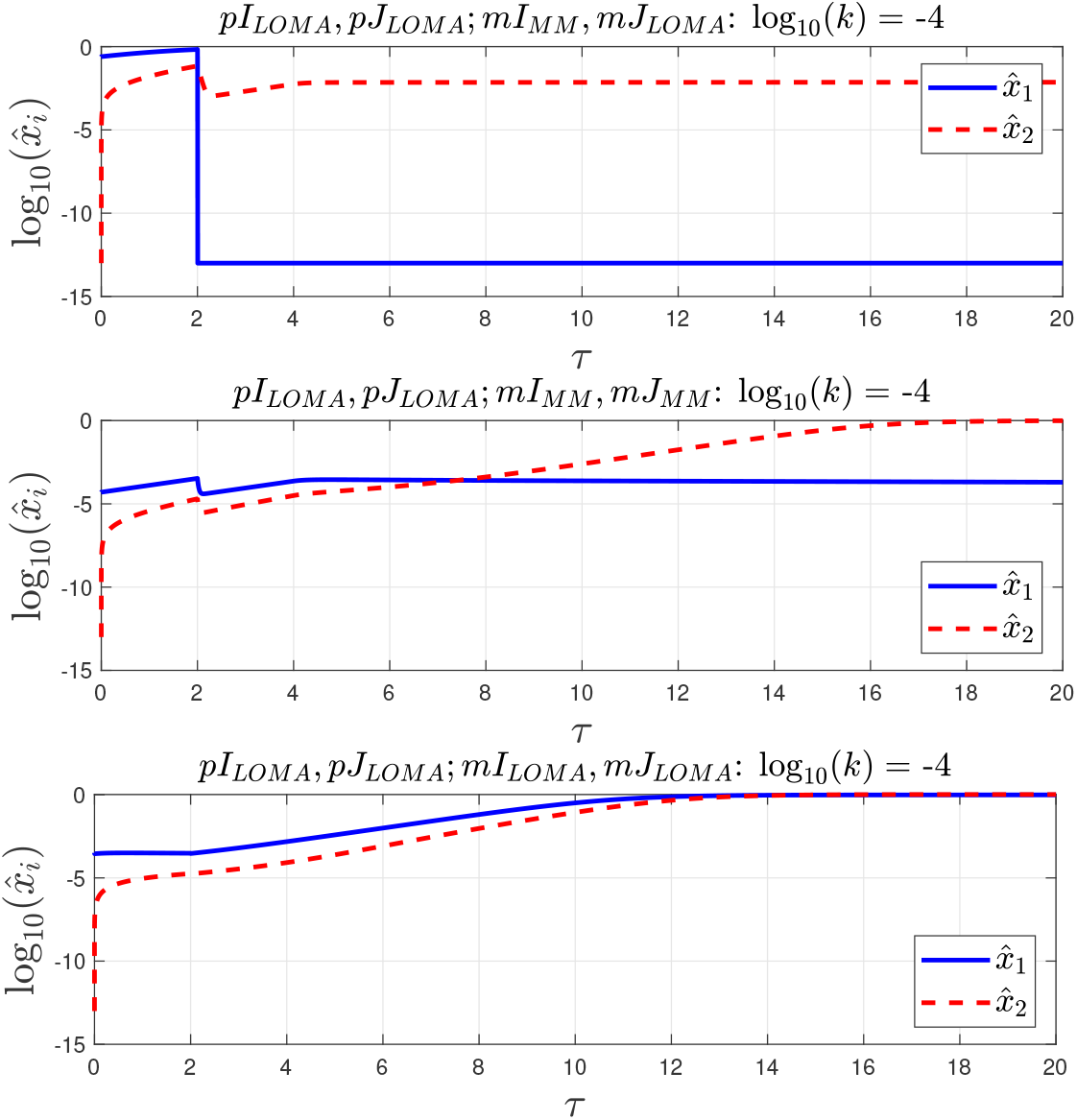
**Top:** Corresponds to Protocol I with 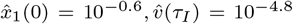 and *I*_*MM*_, *J*_*LOMA*_ secondary tumor phenotype. **Middle:** Corresponds to Protocol I with 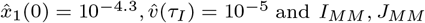 secondary tumor phenotype. **Bottom:** Corresponds to Protocol III with 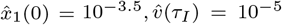 and an initial *I*_*LOMA*_, *J*_*LOMA*_ phenotype for both primary and secondary tumors.

### 5.6 Selected time series

Figures 7-20 represent a large sweep over dosage 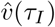 and initial tumor size 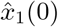, and they provide a visual summary of simulated patient outcomes. However, they can only represent the end-state of the simulations at *τ* = 20, while individual time series are necessary to give a sense of causality and dynamics. Although it is clearly impractical to present time series representing all outcomes summarized in Figures 7-20, we show three particularly illustrative cases below, which coincide with regions in Figures 7-20 where either (i) treatment produced a worse outcome than non-treatment, or (ii) a primary tumor is eliminated but a secondary tumor corresponding to metastasis becomes large.

### 5.7 Comments on simulation studies

In Figures 7-20, we have simulated attempts at treating a tumor four different protocols alongside one untreated baseline. Some of the treatment protocols simulated follow suggestions made elsewhere [31, 33, 34].

Broadly, some of the takeaways from Figures 7-20 are straightforward: for a sufficiently small initial tumor and a sufficiently high treatment dosage, the primary tumor will be eliminated. In the untreated baseline case, primary tumors readily metastasize to a secondary organ above a critical tumor size that the immune system can no longer suppress. For sufficiently high dosages of oncovirus, OVT alone is sufficient to treat even very large tumors, and there are no instances where a primary tumor is eliminated but the metastasis survives or vice-versa.

The simulations also reveal some important nuances. When there is an initial round of immune suppression followed by OVT (Protocol I), and for sufficiently small initial tumors, tumor clearance is greatly enhanced, affirming the suggestions of [31, 33, 34] and consistent with recent clinical trials [77–79]. However, for a sufficiently large initial tumor, the primary tumor can be eliminated even while metastasis to a secondary organ is permitted. This suggests two dangerous possibilities:

- that tumor models based solely on a single tumor can, indeed, falsely predict the success of treatment if metastatic potential is neglected; and
- that the initial round of immune suppression can allow small tumors at the secondary site that would have otherwise been destroyed to grow aggressively.

Across Protocols I and II, where immune suppression is used either before OVT or concurrently with it, small primary tumors 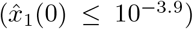 that the immune system would otherwise have suppressed can become viable and large, leading to well-developed metastases. This effect is more pronounced when OVT and immune suppressants are administered at the same time, at *τ* = 0. The danger in this case lies in beginning combination treatment but using too low a dosage of oncovirus. Given the prevalence of severe side effects from OVT, we note that this may have implications for patient survival when treatment must be discontinued; see, for example, discussion in [77–79].

Our inference from the bifurcation diagrams of Sections 4.1-4.2 — that poorly-vascularized tumors may be especially resistant to OVT — is also broadly substantiated. Poorly-vascularized tumors in the secondary organ tend to grow larger, even when the poor vascularization is due to the application of a vascular disrupting agent (VDA). The intuition is reasonably clear: reducing tumor vascularity, all else being equal, tends to shelter the tumor against destruction by the immune system and direct lysis by the administered oncovirus.

## 6 Discussion

### Conclusions

In this paper, we have extended the model of [31, 33] to consider the case of a primary tumor that constantly sheds micrometastases to a nearby secondary organ. We have considered how tumor vascularization — modeled as different functional responses for infectivity of tumor cells *I* and immune killing of tumor cells *J* — affects the bifurcation structure of the resulting six-compartment model, Equations (3). In particular, we find that the bifurcation structure depends rather sensitively on tumor vascularization: well-vascularized tumors are prone to oscillations and limit cycles, while poorly-vascularized tumors have especially long unstable branches along the *k* axis. In the present model, clearance of a large avascular tumor with OVT requires supraphysiological immune killing rates.

We also investigated whether there are conditions under which OVT eliminates a primary tumor only for new, metastatic growth to occur unimpeded at a secondary organ. Comparison of the bifurcation structures for two families of equilibria — those for 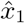 where 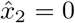, and those for 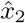 where 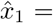 — revealed that such conditions are expected to exist. Poorly vascularized tumors are especially prone to this behavior if allowed to become sufficiently large.

We then studied whether there might exist conditions for which treatment by OVT actively *encourages* metastasis, or a worsening of prognosis compared to an untreated baseline scenario, with special attention to the two-phase protocol suggested by [31, 33, 34]. Under this two-phase protocol, the immune system is deliberately suppressed in order to promote viral buildup, after which immune suppression is reversed, ideally leading to tumor clearance by the combined effects of the oncolytic virus and the replenished immune system. We find that while some tumors may be cleared if the tumor is sufficiently small and the OVT dosage is sufficiently large, there exist dangerous regions of parameter space where OVT and immune modulation clear the primary tumor but may encourage metastasis to the secondary organ.

Informed by our bifurcation analysis in the case of a poorly vascularized tumor, which are especially treatment-resistant, we also investigated whether the use of a vascular disrupting agent (VDA) alongside OVT might worsen simulated patient outcomes. We find that for sufficiently large primary tumors and sufficiently small doses of OVT, the use of a VDA, indeed, drastically worsens the simulated patient outcome, promoting both primary and secondary tumor growth.

### Contribution

The present work, focusing on a six-component nonlinear ODE model, represents a level of analysis that lies in between that of (i) the four-component (or fewer) non-linear ODE models of [2, 27, 31, 33] where precise mathematical statements can be made, and (ii) full, spatiotemporal evolution equations, typically requiring the use of nonlinear partial differential equations, which are difficult to validate and for which a bifurcation analysis of the type made here is intractable. A rigorous, theoretical analysis of the manifold in parameter space separating equilibria that represent positive patient outcomes from negative ones — that is, the separatrix — cannot be made. To our awareness, this is the first model that attempts to develop a bifurcation analysis of a nonlinear ODE model for the treatment of a tumor by OVT that also includes metastasis. We view the present work as a useful bridge between simplified models where exact analysis is possible and more realistic models where analytical or semianalytical results are inaccessible. One consequence of this approach is immediate: the present work shows the existence of conditions for which modeling of the primary tumor alone incorrectly suggests successful treatment.

In our view, the most useful, even if qualitative, contribution of the present work is a warning about the use of immune modulation or vascular disrupting agents alongside OVT. We have found that it is possible for the initial immune suppression phase needed to allow the buildup of therapeutic virus to unintentionally promote metastasis. It is possible for this metastasis to continue unabated even when the primary tumor has been eliminated. We also find that tumor vascularization is expected to play an important role in determining the efficacy of OVT; use of a VDA on a sufficiently large, solid tumor may unintentionally insulate the tumor against the oncolytic virus.

#### Limitations and future work

Apart from the limitations described above, the present work has also made numerous simplifying assumptions. Among them, we have assumed that the shedding rate of the primary tumor to the secondary organ is linear; this is unlikely to be true. However, incorporating a nonlinear shedding rate would have required new assumptions about the saturated shedding rates, which would lie outside of the scope of the present work. We have also assumed that the primary and secondary tumors share parameters, which need not be the case; in principle, a large analysis of the high-dimensional parameter space is possible, but was not immediately relevant to the questions that motivated this work. The assumption of only one additional organ that receives micrometastases from the primary tumor is also oversimplified, but is the minimal necessary step to consider metastasis. Further, we have ignored the cross-seeding of tumor types. We have also considered only a small subset of all possible functional responses; in particular, we have not considered Beddington-DeAngelis or ratio dependent functional responses [27, 46].

It is also somewhat unrealistic to assume, as we have, that the extent of tumor vascularization — and, hence, the functional response — is fixed. Indeed, natural future directions for this work include modeling the evolution of tumor vasculature. Another appealing direction is the development of a full spatiotemporal model that captures the fundamental interactions of immune system, virions, tumor cells and vasculature outside of the simple six-compartment idealization of the present work.

## Declarations

### Ethics approval

Not applicable.

### Consent to participate

Not applicable.

### Consent for publication

Not applicable.

### Availability of data and materials

All simulation data will be made publicly available upon publication.

### Code availability

All code will be made publicly available upon publication.

### Authors’ contribution

TPE and MM conceived of the study idea and contributed to preliminary analysis and numerical experiments. TPE finalized the model and developed the analysis appearing in the final version of the manuscript. TPE wrote the manuscript and both TPE and MM jointly revised it. Both authors have read and approved the final text for submission.

## Appendix A Note on the nondimensionalization

In the main text, we have used a somewhat unorthodox technique to simultaneously nondimensionalize different functional responses *I* and *J* and facilitate a fair comparison between outcomes given a shared set of baseline parameters. Here, we show that the nondimensionalization scheme that we have used properly recovers the intended dimensional quantities. We let

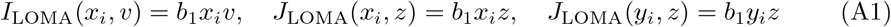

Because *b*_1_ carries units of cell^*™*1^, the physical dimensions of the corresponding rate constants scale as:

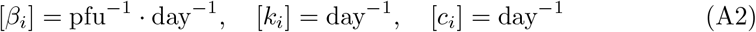

The Michaelis-Menten (MM) forms are

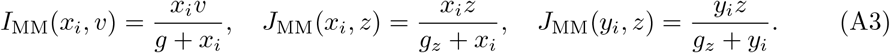

We apply the nondimensionalization defined in Section 3:

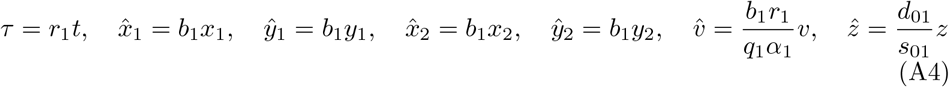

In what follows, we consider the different possible cases for *I* and *J*.

**Case I: *I***_**MM**_, ***J***_**LOMA**_

We assign

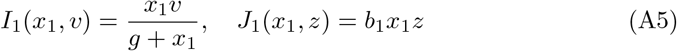

Multiplying the dimensional differential equation for *dx*_1_*/dt* by 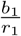and substitutingthe dimensionless variables yields:

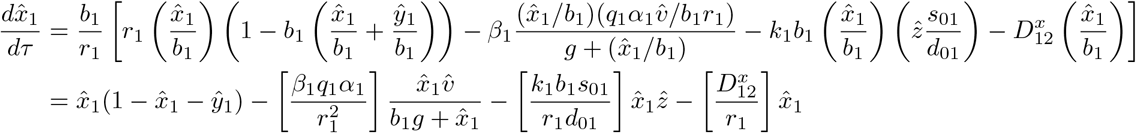

Substituting the baseline parameter groups from Section 3, where 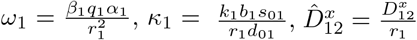, and defining the dimensionless saturation group *G*_*v*_ = *b*_1_*g*, we find

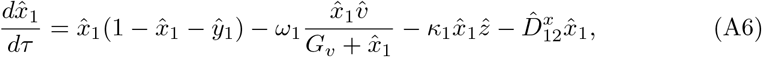

as intended.

**Case II: *I***_**LOMA**_, ***J***_**LOMA**_

We assign

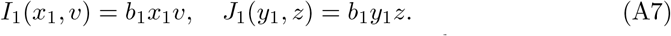

Multiplying the dimensional differential equation for *dy*_1_*/dt* by 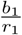yields

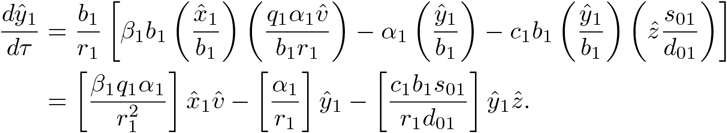

Next, substituting the primary tumor parameter groups 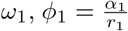, and the baseline immune group 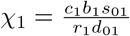, we obtain

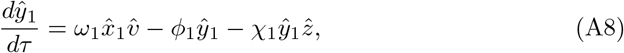

as intended.

**Case III: *I***_**MM**_, ***J***_**MM**_

We assign

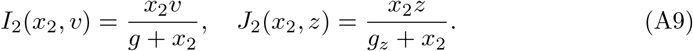

Multiplying the dimensional secondary tumor equation for 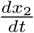by 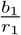yields

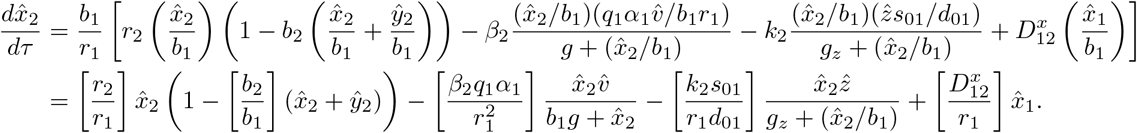

Multiplying the numerator and denominator of the immune saturation term by *b*_1_ leads to

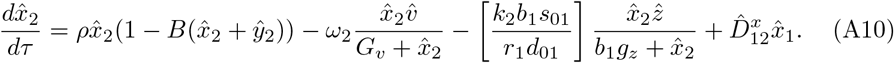

Letting 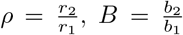, *G*_*v*_ = *b*_1_*g*, and defining the secondary site dimensionless immune saturation constant as *G*_*z*_ = *b*_1_*g*_*z*_, we finally obtain

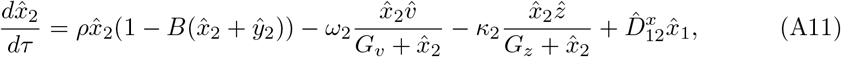

where 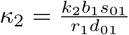 .

## Appendix B Jacobian

The Jacobian **J** ∈ ℝ^6×6^ has non-zero entries

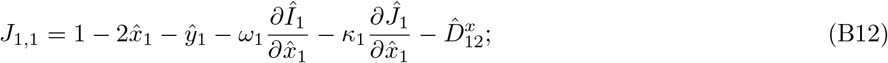

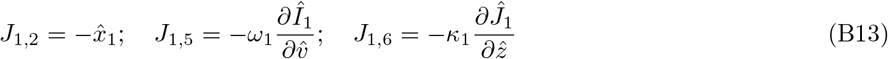

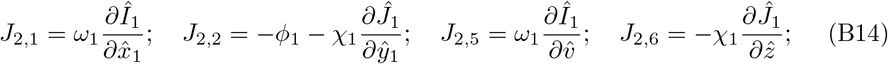

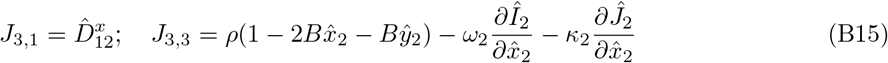

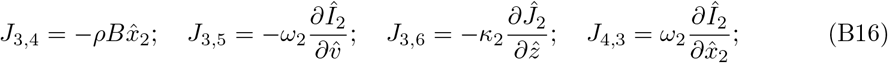

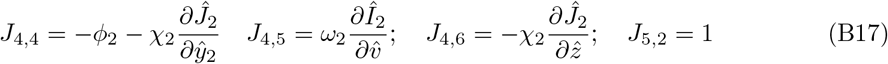

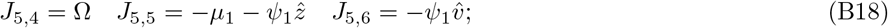

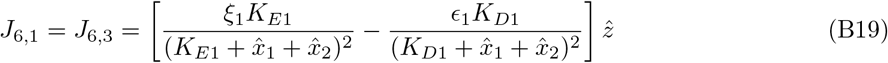

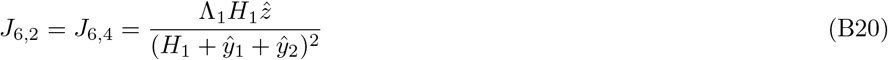

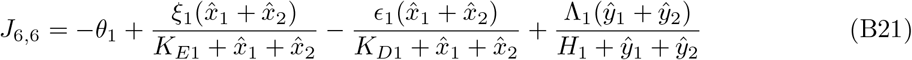

with all other entries of **J** being zero. In general, this matrix appears to have no special structure and we record it here only for completeness.

## References

[1] Martuza, R.L., Malick, A., Markert, J.M., Ruffner, K.L., Coen, D.M.: Experimental therapy of human glioma by means of a genetically engineered virus mutant. Science 252(5007), 854–856 (1991) 10.1126/science.1851332 https://www.science.org/doi/pdf/10.1126/science.1851332

[2] Wodarz, D.: Viruses as antitumor weapons: defining conditions for tumor remission. Cancer Research 61(8), 3501–3507 (2001)

[3] Andtbacka, R.H.I., Collichio, F., Harrington, K.J., Middleton, M.R., Downey, G., hrling, K., Kaufman, H.L.: Final analyses of optim: a randomized phase iii trial of talimogene laherparepvec versus granulocyte-macrophage colony-stimulating factor in unresectable stage iii-iv melanoma. Journal for ImmunoTherapy of Cancer 7(1), 145 (2019) 10.1186/s40425-019-0623-z

[4] Kollmann, C.F., van Montfoort, N., Cordelier, P., Pol, J., Olagnier, D.: Oncolytic virotherapy: Sparking durable anti-tumor immunity through microenvironment modulation. Seminars in Immunology 80, 101994 (2025) 10.1016/j.smim.2025.101994

[5] Hashiro, G., Loh, P.C., Yau, J.T.: The preferential cytotoxicity of reovirus for certain transformed cell lines. Archives of Virology 54(4), 307–315 (1977) 10.1007/BF01314776

[6] Coffey, M.C., Strong, J.E., Forsyth, P.A., Lee, P.W.K.: Reovirus therapy of tumors with activated ras pathway. Science 282(5392), 1332–1334 (1998) 10.1126/science.282.5392.1332 https://www.science.org/doi/pdf/10.1126/science.282.5392.1332

[7] Stojdl, D.F., Lichty, B., Knowles, S., Marius, R., Atkins, H., Sonenberg, N., Bell, J.C.: Exploiting tumor-specific defects in the interferon pathway with a previously unknown oncolytic virus. Nature Medicine 6(7), 821–825 (2000) 10.1038/77558

[8] Heise, C., Hermiston, T., Johnson, L., Brooks, G., Sampson-Johannes, A., Williams, A., Hawkins, L., Kirn, D.: An adenovirus e1a mutant that demonstrates potent and selective systemic anti-tumoral efficacy. Nature Medicine 6(10), 1134–1139 (2000) 10.1038/80474

[9] Wong, R.J., Joe, J.K., Kim, S.-H., Shah, J.P., Horsburgh, B., Fong, Y.: Oncolytic herpesvirus effectively treats murine squamous cell carcinoma and spreads by natural lymphatics to treat sites of lymphatic metastases. Human Gene Therapy 13(10), 1213–1223 (2002) 10.1089/104303402320138998 https://journals.sagepub.com/doi/pdf/10.1089/104303402320138998. PMID: 12133274

[10] Sharon, D., Schümann, M., MacLeod, S., McPherson, R., Chaurasiya, S., Shaw, A., Hitt, M.M.: 2-aminopurine enhances the oncolytic activity of an e1b-deleted adenovirus in hepatocellular carcinoma cells. PLOS ONE 8(6), 1–14 (2013) 10.1371/journal.pone.0065222

[11] Bhat, R., Dempe, S., Dinsart, C., Rommelaere, J.: Enhancement of nk cell antitumor responses using an oncolytic parvovirus. International Journal of Cancer 128(4), 908–919 (2011) 10.1002/ijc.25415 https://onlinelibrary.wiley.com/doi/pdf/10.1002/ijc.25415

[12] Ogbomo, H., Zemp, F.J., Lun, X., Zhang, J., Stack, D., Rahman, M.M., Mcfadden, G., Mody, C.H., Forsyth, P.A.: Myxoma virus infection promotes nk lysis of malignant gliomas in vitro and in vivo. PLOS ONE 8(6), 1–14 (2013) 10.1371/journal.pone.0066825

[13] Munn, D.H., Bronte, V.: Immune suppressive mechanisms in the tumor microenvironment. Current Opinion in Immunology 39, 1–6 (2016) 10.1016/j.coi.2015.10.009. Lymphocyte development and activation * Tumour immunology

[14] Labani-Motlagh, A., Ashja-Mahdavi, M., Loskog, A.: The tumor microenvironment: A milieu hindering and obstructing antitumor immune responses. Frontiers in Immunology Volume 11-2020 (2020) 10.3389/fimmu.2020.00940

[15] Tufail, M., Jiang, C.-H., Li, N.: Immune evasion in cancer: mechanisms and cutting-edge therapeutic approaches. Signal Transduction and Targeted Therapy 10(1), 227 (2025) 10.1038/s41392-025-02280-1

[16] Chaurasiya, S., Chen, N.G., Fong, Y.: Oncolytic viruses and immunity. Current Opinion in Immunology 51, 83–90 (2018) 10.1016/j.coi.2018.03.008

[17] Liu, T.-C., Galanis, E., Kirn, D.: Clinical trial results with oncolytic virotherapy: a century of promise, a decade of progress. Nature Clinical Practice Oncology 4(2), 101–117 (2007) 10.1038/ncponc0736

[18] Aurelian, L.: Oncolytic viruses as immunotherapy: progress and remaining challenges. OncoTargets and Therapy 9, 2627–2637 (2016) 10.2147/OTT.S63049

[19] Alam, S.M.S., Tasnim, M., Amin, T., Islam, S.M.B.U.: A new era in oncology: Clinical insights into the application of oncolytic viruses. Cancer Control 32, 10732748251382295 (2025) 10.1177/10732748251382295 https://doi.org/10.1177/10732748251382295

[20] Prestwich, R.J., Errington, F., Diaz, R.M., Pandha, H.S., Harrington, K.J., Melcher, A.A., Vile, R.G.: The case of oncolytic viruses versus the immune system: Waiting on the judgment of solomon. Human Gene Therapy 20(10), 1119–1132 (2009) 10.1089/hum.2009.135 https://journals.sagepub.com/doi/pdf/10.1089/hum.2009.135. PMID: 19630549

[21] Wu, J.T., Kirn, D.H., Wein, L.M.: Analysis of a three-way race between tumor growth, a replication-competent virus and an immune response. Bulletin of Mathematical Biology 66(4), 605–625 (2004) 10.1016/j.bulm.2003.08.016

[22] Yin, A., Moes, D.J.A.R., Hasselt, J.G.C., Swen, J.J., Guchelaar, H.-J.: A review of mathematical models for tumor dynamics and treatment resistance evolution of solid tumors. CPT: Pharmacometrics & Systems Pharmacology 8(10), 720–737 (2019) 10.1002/psp4.12450 https://ascpt.onlinelibrary.wiley.com/doi/pdf/10.1002/psp4.12450

[23] Guo, E., Dobrovolny, H.M.: Mathematical modeling of oncolytic virus therapy reveals role of the immune response. Viruses 15(9) (2023) 10.3390/v15091812

[24] Kuznetsov, V.A., Makalkin, I.A., Taylor, M.A., Perelson, A.S.: Nonlinear dynamics of immunogenic tumors: Parameter estimation and global bifurcation analysis. Bulletin of Mathematical Biology 56(2), 295–321 (1994) 10.1016/S0092-8240(05)80260-5

[25] Fukuhara, H., Ino, Y., Todo, T.: Oncolytic virus therapy: A new era of cancer treatment at dawn. Cancer Science 107(10), 1373–1379 (2016) 10.1111/cas.13027 https://onlinelibrary.wiley.com/doi/pdf/10.1111/cas.13027

[26] Mahasa, K.J., Eladdadi, A., Pillis, L., Ouifki, R.: Oncolytic potency and reduced virus tumor-specificity in oncolytic virotherapy. a mathematical modelling approach. PLOS ONE 12(9), 1–25 (2017) 10.1371/journal.pone.0184347

[27] Wei, H.-C.: A mathematical model of tumour growth with beddington–deangelis functional response: a case of cancer without disease. Journal of Biological Dynamics 12(1), 194–210 (2018) 10.1080/17513758.2017.1418028 https://doi.org/10.1080/17513758.2017.1418028. PMID: 29322865

[28] Garcia, V., Bonhoeffer, S., Fu, F.: Cancer-induced immunosuppression can enable effectiveness of immunotherapy through bistability generation: A mathematical and computational examination. Journal of Theoretical Biology 492, 110185 (2020) 10.1016/j.jtbi.2020.110185

[29] Sardar, M., Khajanchi, S., Biswas, S., Abdelwahab, S.F., Nisar, K.S.: Exploring the dynamics of a tumor-immune interplay with time delay. Alexandria Engineering Journal 60(5), 4875–4888 (2021) 10.1016/j.aej.2021.03.041

[30] Mahasa, K.J., Ouifki, R., Eladdadi, A., Pillis, L.: A combination therapy of oncolytic viruses and chimeric antigen receptor t cells: a mathematical model proof-of-concept. Mathematical Biosciences and Engineering 19(5), 4429–4457 (2022) 10.3934/mbe.2022205

[31] Vithanage, G.V.R.K., Wei, H.-C., Jang, S.R.-J.: The role of tumor activation and inhibition with saturation effects in a mathematical model of tumor and immune system interactions undergoing oncolytic viral therapy. Mathematical Methods in the Applied Sciences 46(9), 10787–10813 (2023) 10.1002/mma.9152 https://onlinelibrary.wiley.com/doi/pdf/10.1002/mma.9152

[32] Mohammadnejad, N., Hillen, T.: Qualitative optimization of oncolytic virotherapy and immune therapy combination treatments. Mathematical Biosciences and Engineering 23(2), 388–420 (2026) 10.3934/mbe.2026016

[33] Vithanage, G.V.R.K., Wei, H.-C., Jang, S.R.-J.: Bistability in a model of tumorimmune system interactions with an oncolytic viral therapy. Mathematical Biosciences and Engineering 19(2), 1559–1587 (2022) 10.3934/mbe.2022072

[34] Rojas, J.J., Sampath, P., Hou, W., Thorne, S.H.: Defining effective combinations of immune checkpoint blockade and oncolytic virotherapy. Clinical Cancer Research 21(24), 5543–5551 (2015) 10.1158/1078-0432.CCR-14-2009https://aacrjournals.org/clincancerres/article-pdf/21/24/5543/2029348/5543.pdf

[35] Hellman, S., Weichselbaum, R.R.: Oligometastases. Journal of Clinical Oncology 13(1), 8–10 (1995) 10.1200/JCO.1995.13.1.8 https://ascopubs.org/doi/pdf/10.1200/JCO.1995.13.1.8. PMID: 7799047

[36] AlGhamdi, H., Dhont, J., Krayem, M., De Bruyn, P., Engels, B., Van Gestel, D., Begin, R.: The road to dissemination: The concept of oligometastases and the barriers for widespread disease. Cancers 14(8) (2022) 10.3390/cancers14082046

[37] Seyfried, T.N., Huysentruyt, L.C.: On the origin of cancer metastasis. Critical Reviews in Oncogenesis 18(1-2), 43–73 (2013) 10.1615/CritRevOncog.v18.i1-2.40

[38] Nataraj, N.B., Marrocco, I., Yarden, Y.: Roles for growth factors and mutations in metastatic dissemination. Biochemical Society Transactions 49(3), 1409–1423 (2021) 10.1042/BST20210048 https://portlandpress.com/biochemsoctrans/article-pdf/49/3/1409/916204/bst-2021-0048c.pdf

[39] Teng, M.W.L., Swann, J.B., Koebel, C.M., Schreiber, R.D., Smyth, M.J.: Immune-mediated dormancy: an equilibrium with cancer. Journal of Leukocyte Biology 84(4), 988–993 (2008) 10.1189/jlb.1107774 https://jlb.onlinelibrary.wiley.com/doi/pdf/10.1189/jlb.1107774

[40] Payne, K.K., Keim, R.C., Graham, L., Idowu, M.O., Wan, W., Wang, X.-Y., Toor, A.A., Bear, H.D., Manjili, M.H.: Tumor-reactive immune cells protect against metastatic tumor and induce immunoediting of indolent but not quiescent tumor cells. Journal of Leukocyte Biology 100(3), 625–635 (2016) 10.1189/jlb.5A1215-580R https://jlb.onlinelibrary.wiley.com/doi/pdf/10.1189/jlb.5A1215-580R

[41] Bridle, B.W., Stephenson, K.B., Boudreau, J.E., Koshy, S., Kazdhan, N., Pullenayegum, E., Brunelliére, J., Bramson, J.L., Lichty, B.D., Wan, Y.: Potentiating cancer immunotherapy using an oncolytic virus. Molecular Therapy 18(8), 1430–1439 (2010) 10.1038/mt.2010.98

[42] Jain, R.K.: Delivery of molecular and cellular medicine to solid tumors1pii of original article: S0169-409x(97)00027-6. the article was originally published in advanced drug delivery reviews 26 (1997) 71–90.1. Advanced Drug Delivery Reviews 46(1), 149–168 (2001) 10.1016/S0169-409X(00)00131-9Special issue dedicated to Dr. Eric Tomlinson, Advanced Drug Delivery Reviews, A Selection of the Most Highly Cited Articles, 1991–1998

[43] Marchini, A., Scott, E.M., Rommelaere, J.: Overcoming barriers in oncolytic virotherapy with hdac inhibitors and immune checkpoint blockade. Viruses 8(1) (2016) 10.3390/v8010009

[44] Luzzi, K.J., MacDonald, I.C., Schmidt, E.E., Kerkvliet, N., Morris, V.L., Chambers, A.F., Groom, A.C.: Multistep nature of metastatic inefficiency: Dormancy of solitary cells after successful extravasation and limited survival of early micrometastases. The American Journal of Pathology 153(3), 865–873 (1998) 10.1016/S0002-9440(10)65628-3

[45] Tabu, K., Mawatari, S., Oda, K., Kumagai, K., Inada, Y., Uto, H., Saisyoji, A., Hiramine, Y., Hashiguchi, M., Tamai, T., Hori, T., Fujisaki, K., Imanaka, D., Kure, T., Taniyama, O., Toyodome, A., Ijuin, S., Sakae, H., Sakurai, K., Moriuchi, A., Kanmura, S., Ido, A.: Hypovascular tumors developed into hepatocellular carcinoma at a high rate despite the elimination of hepatitis c virus by direct-acting antivirals. PLOS ONE 15(8), 1–14 (2020) 10.1371/journal.pone.0237475

[46] Radunskaya, A., Sack, J.: Kill rates by immune cells: Ratio-dependent, or mass action? Journal of Theoretical Biology 582, 111748 (2024) 10.1016/j.jtbi.2024.111748

[47] Liotta, L.A., Kleinerman, J., Saidel, G.M.: Quantitative relationships of intravascular tumor cells, tumor vessels, and pulmonary metastases following tumor implantation. Cancer Research 34(5), 997–1004 (1974) https://aacrjournals.org/cancerres/article-pdf/34/5/997/2392400/cr0340050997.pdf

[48] Avanzini, S., Kurtz, D.M., Chabon, J.J., Moding, E.J., Hori, S.S., Gambhir, S.S., Alizadeh, A.A., Diehn, M., Reiter, J.G.: A mathematical model of ctdna shedding predicts tumor detection size. Science Advances 6(50), 4308 (2020) 10.1126/sciadv.abc4308 https://www.science.org/doi/pdf/10.1126/sciadv.abc4308

[49] Massagué, J., Obenauf, A.C.: Metastatic colonization by circulating tumour cells. Nature 529(7586), 298–306 (2016) 10.1038/nature17038

[50] Thiele, J.-A., Bethel, K., Králíčková, M., Kuhn, P.: Circulating tumor cells: Fluid surrogates of solid tumors. Annual Review of Pathology: Mechanisms of Disease 12(Volume 12, 2017), 419–447 (2017) 10.1146/annurev-pathol-052016-100256

[51] Szczurek, E., Krüger, T., Klink, B., Beerenwinkel, N.: A mathematical model of the metastatic bottleneck predicts patient outcome and response to cancer treatment. PLOS Computational Biology 16(10), 1–20 (2020) 10.1371/journal.pcbi.1008056

[52] Stoletov, K., Beatty, P.H., Lewis, J.D.: Novel therapeutic targets for cancer metastasis. Expert Review of Anticancer Therapy 20(2), 97–109 (2020) 10.1080/14737140.2020.1718496 https://doi.org/10.1080/14737140.2020.1718496. PMID: 31997674

[53] Zhang, C., Liu, Z., Li, X., Li, S., Tan, Y.: Mechano-regulation of tumor metastasis: from intravasation to extravasation. Med-X 4(1), 20 (2026) 10.1007/s44258-026-00091-9

[54] Wang, W.-C., Zhang, X.-F., Peng, J., Li, X.-F., Wang, A.-L., Bie, Y.-Q., Shi, L.-H., Lin, M.-B., Zhang, X.-F.: Survival mechanisms and influence factors of circulating tumor cells. BioMed Research International 2018(1), 6304701 (2018) 10.1155/2018/6304701 https://onlinelibrary.wiley.com/doi/pdf/10.1155/2018/6304701

[55] Paresishvili, T., Kakabadze, Z.: Challenges and opportunities associated with drug delivery for the treatment of solid tumors. Oncology Reviews 17, 10577 (2023) 10.3389/or.2023.10577

[56] Padera, T.P., Stoll, B.R., Tooredman, J.B., Capen, D., Tomaso, E.d., Jain, R.K.: Cancer cells compress intratumour vessels. Nature 427(6976), 695–695 (2004) 10.1038/427695a

[57] Stylianopoulos, T., Martin, J.D., Chauhan, V.P., Jain, S.R., Diop-Frimpong, B., Bardeesy, N., Smith, B.L., Ferrone, C.R., Hornicek, F.J., Boucher, Y., Munn, L.L., Jain, R.K.: Causes, consequences, and remedies for growth-induced solid stress in murine and human tumors. Proceedings of the National Academy of Sciences 109(38), 15101–15108 (2012) 10.1073/pnas.1213353109 https://www.pnas.org/doi/pdf/10.1073/pnas.1213353109

[58] Jain, R.K., Martin, J.D., Stylianopoulos, T.: The role of mechanical forces in tumor growth and therapy. Annual Review of Biomedical Engineering 16(Volume 16, 2014), 321–346 (2014) 10.1146/annurev-bioeng-071813-105259

[59] Nia, H.T., Munn, L.L., Jain, R.K.: Physical traits of cancer. Science 370(6516), 0868 (2020) 10.1126/science.aaz0868 https://www.science.org/doi/pdf/10.1126/science.aaz0868

[60] Siemann, D.W.: The unique characteristics of tumor vasculature and preclinical evidence for its selective disruption by tumor-vascular disrupting agents. Cancer Treatment Reviews 37(1), 63–74 (2011) 10.1016/j.ctrv.2010.05.001

[61] Jarrett, A.M., Lima, E.A.B.F., II, D.A.H., McKenna, M.T., Feng, X., Ekrut, D.A., Resende, A.C.M., Brock, A., Yankeelov, T.E.: Mathematical models of tumor cell proliferation: A review of the literature. Expert Review of Anticancer Therapy 18(12), 1271–1286 (2018) 10.1080/14737140.2018.1527689 https://doi.org/10.1080/14737140.2018.1527689. PMID: 30252552

[62] Leung, C.T., Brugge, J.S.: Tumor self-seeding: Bidirectional flow of tumor cells. Cell 139(7), 1226–1228 (2009) 10.1016/j.cell.2009.12.013

[63] Kim, M.-Y., Oskarsson, T., Acharyya, S., Nguyen, D.X., Zhang, X.H.-F., Norton, L., Massagué, J.: Tumor self-seeding by circulating cancer cells. Cell 139(7), 1315–1326 (2009) 10.1016/j.cell.2009.11.025

[64] Aguirre-Ghiso, J.A.: On the theory of tumor self-seeding: implications for metastasis progression in humans. Breast Cancer Research 12(2), 304 (2010)10.1186/bcr2561

[65] Scott, J.G., Basanta, D., Anderson, A.R.A., Gerlee, P.: A mathematical model of tumour self-seeding reveals secondary metastatic deposits as drivers of primary tumour growth. Journal of The Royal Society Interface 10(82), 20130011 (2013) 10.1098/rsif.2013.0011 https://royalsocietypublishing.org/rsif/article-df/doi/10.1098/rsif.2013.0011/914499/rsif.2013.0011.pdf

[66] Kipnis, P., Ramanathan, D., Hoehn, R., Jethwa, A.R., Karakla, D.W., Rohr, B., Sutter, C.M., Mark, J.R., Khaja, S.F., Li, S., Thuener, J., Carroll, B.T.: Tumor seeding across specialties: a systematic review. Frontiers in Oncology Volume 14-2024 (2024) 10.3389/fonc.2024.1464767

[67] Mañes, S., Gómez-Moutón, C., Lacalle, R.A., Jiménez-Baranda, S., Mira, E., Martínez-A, C.: Mastering time and space: immune cell polarization and chemo-taxis. Seminars in Immunology 17(1), 77–86 (2005) 10.1016/j.smim.2004.09.005. Spacial Organization in Immune Cell Signaling

[68] Jin, T., Xu, X., Hereld, D.: Chemotaxis, chemokine receptors and human disease. Cytokine 44(1), 1–8 (2008) 10.1016/j.cyto.2008.06.017

[69] Zhou, J., Xiang, Y., Yoshimura, T., Chen, K., Gong, W., Huang, J., Zhou, Y., Yao, X., Bian, X., Wang, J.M.: The role of chemoattractant receptors in shaping the tumor microenvironment. BioMed Research International 2014(1), 751392 (2014) 10.1155/2014/751392 https://onlinelibrary.wiley.com/doi/pdf/10.1155/2014/751392

[70] Ryan, A.T., Kim, M., Lim, K.: Immune cell migration to cancer. Cells 13(10) (2024) 10.3390/cells13100844

[71] Koujima, T., Tazawa, H., Ieda, T., Araki, H., Fushimi, T., Shoji, R., Kuroda, S., Kikuchi, S., Yoshida, R., Umeda, Y., Teraishi, F., Urata, Y., Mizuguchi, H., Fujiwara, T.: Oncolytic virus-mediated targeting of the erk signaling pathway inhibits invasive propensity in human pancreatic cancer. Molecular Therapy-Oncolytics 17, 107–117 (2020) 10.1016/j.omto.2020.03.016

[72] Butler, T.P., Gullino, P.M.: Quantitation of cell shedding into efferent blood of mammary adenocarcinoma. Cancer Research 35(3), 512–516 (1975)

[73] Swartz, M.A., Kristensen, C.A., Melder, R.J., Roberge, S., Calautti, E., Fukumura, D., Jain, R.K.: Cells shed from tumours show reduced clonogenicity, resistance to apoptosis, and in vivo tumorigenicity. British Journal of Cancer 81(5), 756–759 (1999) 10.1038/sj.bjc.6690760

[74] Nowell, P.C.: The clonal evolution of tumor cell populations. Science 194(4260), 23–28 (1976) 10.1126/science.959840 https://www.science.org/doi/pdf/10.1126/science.959840

[75] Sylvester, B.E., Vakiani, E.: Tumor evolution and intratumor heterogeneity in colorectal carcinoma: insights from comparative genomic profiling of primary tumors and matched metastases. Journal of Gastrointestinal Oncology 6(6) (2015)

[76] Dang, H.X., Krasnick, B.A., White, B.S., Grossman, J.G., Strand, M.S., Zhang, J., Cabanski, C.R., Miller, C.A., Fulton, R.S., Goedegebuure, S.P., Fronick, C.C., Griffith, M., Larson, D.E., Goetz, B.D., Walker, J.R., Hawkins, W.G., Strasberg, S.M., Linehan, D.C., Lim, K.H., Lockhart, A.C., Mardis, E.R., Wilson, R.K., Ley, T.J., Maher, C.A., Fields, R.C.: The clonal evolution of metastatic colorectal cancer. Science Advances 6(24), 9691 (2020) 10.1126/sciadv.aay9691 https://www.science.org/doi/pdf/10.1126/sciadv.aay9691

[77] Cerullo, V., Diaconu, I., Kangasniemi, L., Rajecki, M., Escutenaire, S., Koski, A., Romano, V., Rouvinen, N., Tuuminen, T., Laasonen, L., Partanen, K., Kauppinen, S., Joensuu, T., Oksanen, M., Holm, S.-L., Haavisto, E., Karioja-Kallio, A., Kanerva, A., Pesonen, S., Arstila, P.T., Hemminki, A.: Immunological effects of low-dose cyclophosphamide in cancer patients treated with oncolytic adenovirus. Molecular Therapy 19(9), 1737–1746 (2011) 10.1038/mt.2011.113

[78] Roulstone, V., Khan, K., Pandha, H.S., Rudman, S., Coffey, M., Gill, G.M., Melcher, A.A., Vile, R., Harrington, K.J., Bono, J., Spicer, J.: Phase i trial of cyclophosphamide as an immune modulator for optimizing oncolytic reovirus delivery to solid tumors. Clinical Cancer Research 21(6), 1305–1312 (2015) 10.1158/1078-0432.CCR-14-1770 https://aacrjournals.org/clincancerres/article-pdf/21/6/1305/2029124/1305.pdf

[79] Toulmonde, M., Cousin, S., Kind, M., Guegan, J.-P., Bessede, A., Le Loarer, F., Perret, R., Cantarel, C., Bellera, C., Italiano, A.: Randomized phase 2 trial of intravenous oncolytic virus jx-594 combined with low-dose cyclophosphamide in patients with advanced soft-tissue sarcoma. Journal of Hematology & Oncology 15(1), 149 (2022) 10.1186/s13045-022-01370-9

[80] Chaurasiya, S., Fong, Y., Warner, S.G.: Oncolytic virotherapy for cancer: Clinical experience. Biomedicines 9(4) (2021) 10.3390/biomedicines9040419

[81] Fowler, A.C.: Atto-foxes and other minutiae. Bulletin of Mathematical Biology 83(10), 104 (2021) 10.1007/s11538-021-00936-x

